# Two functionally distinct HEATR5 protein complexes are defined by fast-evolving co-factors in yeast

**DOI:** 10.1101/2023.08.24.554671

**Authors:** Lucas J. Marmorale, Huan Jin, Thomas G. Reidy, Brandon Palomino-Alonso, Christopher Zysnarski, Fatima Jordan-Javed, Sagar Lahiri, Mara C Duncan

## Abstract

The highly conserved HEATR5 proteins are best known for their roles in membrane traffic mediated by the adaptor protein complex-1 (AP1). HEATR5 proteins rely on fast-evolving co-factors to bind to AP1. However, how HEATR5 proteins interact with these co-factors is unknown. Here, we report that the budding yeast HEATR5 protein, Laa1, functions in two biochemically distinct complexes. These complexes are defined by a pair of mutually exclusive Laa1-binding proteins, Laa2 and the previously uncharacterized Lft1/Yml037c. Despite limited sequence similarity, biochemical analysis and structure predictions indicate that Lft1 and Laa2 bind Laa1 via structurally similar mechanisms. Both Laa1 complexes function in intra-Golgi recycling. However, only the Laa2-Laa1 complex binds to AP1 and contributes to its localization. Finally, structure predictions indicate that human HEATR5 proteins bind to a pair of fast-evolving interacting partners via a mechanism similar to that observed in yeast. These results reveal mechanistic insight into how HEATR5 proteins bind their co-factors and indicate that Laa1 performs functions besides recruiting AP1.

## Introduction

Membrane trafficking impacts nearly every aspect of cell physiology by regulating the proteomes of numerous organelles, including the Golgi, plasma membrane, endosomes, and lysosomes. Many trafficking pathways rely on soluble proteins that assemble into a proteinaceous coat on the cytosolic surface of membranes. The coat selects protein cargo, deforms the membrane, and ultimately pinches the membrane off into a spherical, tubular, or more complicated transport carrier. Consistent with this fundamental role of coat proteins in maintaining organelle function, mutations in coat proteins can cause devastating human diseases (Duncan, 2022; Sanger et al., 2019; Shin et al., 2021).

Despite decades of research, many coat components remain poorly characterized, including the highly conserved HEATR5 protein family. This family includes Laa1 in budding yeast, Sip1 in fission yeast, SOAP-1 in worms, HEATR5B in flies (also known as CG2747), HEATR5B and HEATR5A in mammals (also known as p200a and p200b), and Sweetie in plants (Fernandez and Payne, 2006; Gillard et al., 2015; Le Bras et al., 2012; Yu et al., 2012). HEATR5 is essential in flies, worms, and mice, and mutations in HEATR5B cause a neurological syndrome characterized by pontocerebellar hypoplasia, neonatal seizures, severe intellectual disability, and motor delay (Ghosh et al., 2021; Gillard et al., 2015; Le Bras et al., 2012). However, our knowledge about the functions of HEATR5 proteins and their molecular mechanisms is limited.

HEATR5 proteins localize to the trans-Golgi network and endosomes and, until now, were mainly thought to promote AP1 localization to membranes. HEATR5 proteins bind to AP1 indirectly via one or more fast-evolving co-factors (Kuznetsov et al., 2023). In budding yeast, Laa1 forms a complex with the co- factor Laa2, which mediates the interaction with AP1 (Zysnarski et al., 2019). In humans, HEATR5b simultaneously binds two co-factors, Aftiphilin and γ-Synergin, which interact directly with AP1 and are thought to mediate its interaction with HEATR5b (Hirst et al., 2005). However, it remains unknown how HEATR5 proteins bind these critical co-factors and whether HEATR5 proteins perform functions in addition to promoting AP1 membrane localization.

In the present study, we sought to clarify the molecular mechanisms of the yeast HEATR5 protein Laa1 by identifying new co-factors. We identified a previously uncharacterized protein, Yml037c, as a novel Laa1 co-factor. We refer to Yml037c as Lft1 for ***L****aa1 **F**unction **T**wo 1.* We defined how Lft1 and Laa2 bind Laa1 and revealed that they form mutually exclusive complexes with Laa1. In addition, we showed that the Lft1-Laa1 complex does not bind to or contribute to AP1 localization. These findings suggest that Laa1 has functions besides promoting AP1 membrane localization. Finally, we determined that human HEATR5 proteins likely use an interface similar to that found in yeast to bind to a pair of mutually exclusive co-factors. Together, these findings indicate that HEATR5 proteins likely play multiple roles in eukaryotic cells.

### Identification of Lft1

Laa1 is a large protein with the potential to make many physical interactions. However, it is only known to interact directly with Laa2. To identify additional co-factors, we performed immunoprecipitation followed by mass spectrometry analysis. In the Laa1 samples, Laa2, subunits of AP1, and several additional proteins were identified but not in control samples (the entire mass spectrometry data is available in Table S1). We hypothesized that if any of these additional proteins represented co-factors of Laa1, a deletion of its gene would have similar phenotypes to the deletion of *LAA1*. Therefore, we examined the HOP-profile dataset, which contains information about the growth effect of ∼1800 different chemical treatments on gene deletion strains in the yeast deletion collection (Hoepfner et al., 2014). We looked for gene deletions whose sensitivity and resistance profiles correlate with those of *laa1Δ*. Among the top twenty-five correlated genes, we found a single uncharacterized gene, *YML037c,* whose gene product was also identified in our mass spectrometry samples. Based on the results described below, we refer to Yml037c as Lft1 (*Laa1 Function Two 1*).

To determine whether Lft1 interacted with Laa1, we tagged Lft1 with the FLAG epitope at its endogenous locus and performed co-immunoprecipitation with Laa1 tagged with the MYC epitope at its endogenous locus. We found that Lft1 interacted with Laa1 (Fig. 1A). We next asked whether Lft1 localization depends on Laa1. We tagged Lft1 with GFP at its endogenous locus and compared its localization in wild- type and cells lacking Laa1. In wild-type cells, Lft1-GFP localizes to one or more puncta per cell. In cells lacking Laa1, this localization is abolished (Fig. 1B, C). Because Lft1-GFP puncta were completely undetectable, we sought to determine if Laa1 altered Lft1 abundance. To do this, we monitored Lft1-HA abundance in cells lacking Laa1 by immunoblot analysis. Lft1-HA abundance was dramatically lower in cells lacking Laa1 (Fig. 1D, E). This lower abundance suggests that Lft1 is destabilized in the absence of Laa1 and indicates that Lft1 forms a complex with Laa1.

**Figure 1.**
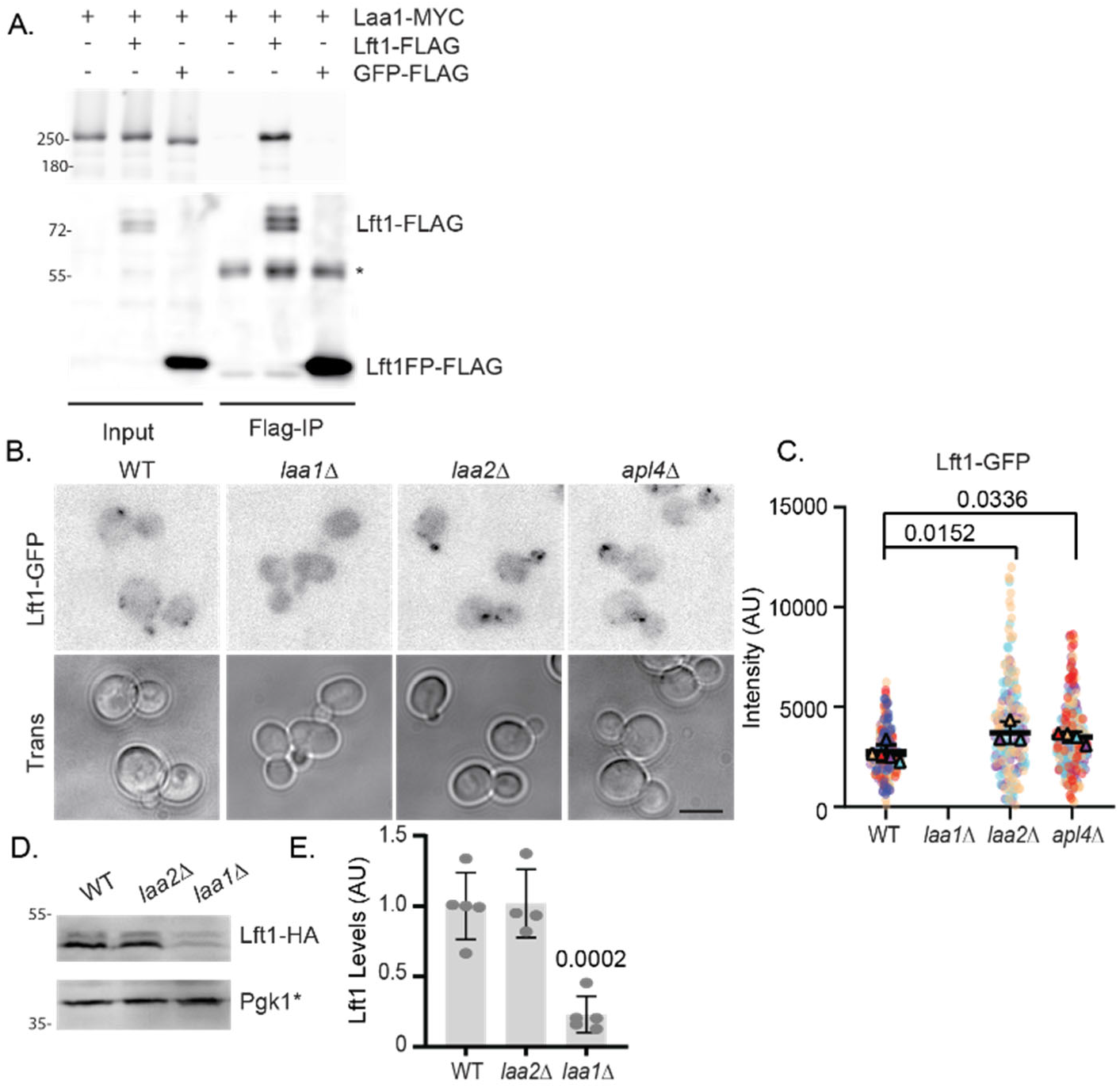
Lft1 binds Laa1 and depends on Laa1 for protein stability. A. Lft1 binds Laa1 as detected by immunoprecipitation. Flag-tagged proteins were immunoprecipitated from cells expressing indicated gene fusions expressed from their endogenous loci and probed for Flag and HA. * marks antibody band B. Lft1-GFP puncta are not detectable in cells lacking Laa1 and are slightly more intense in cells lacking Laa2 or AP1. Cells with indicated genotypes were imaged as described in materials and methods. Scale bar is 5 μm C. Quantification of Lft1 puncta intensity in indicated cells. Charts show super-plots of median values from replicate experiments plotted over individual intensity measurements which are color coded for each replicate experiment. The standard deviations of the median values are shown. P-values derive from Dunnett’s multiple comparisons test of the mean values of replicate experiments D. Lft1 abundance is lower in cells lacking Laa1 but not Laa2. Lysates of cells of indicated genotypes were probed for HA or Pgk1*, a breakdown product of Pgk1 used as a lysis control. E. Charts of intensity values from D. The standard deviations are shown. P-values derive from Dunnett’s multiple comparisons test. Chart shows standard deviations.

### Lft1 and Laa2 are mutually exclusive for binding to Laa1

Because Laa1 is known to form a complex with Laa2 and Laa2 is important for Laa1 localization (Zysnarski et al., 2019), we asked whether loss of Laa2 disrupts Lft1 localization. In contrast to the effect of Laa1, we found that the loss of Laa2 did not disrupt Lft1 localization. Instead, Lft1 puncta were slightly brighter than in wild-type cells (Fig. 1B, C). In addition, Lft1 abundance was unaffected by the loss of Laa2 as detected by immunoblot analysis (Fig. 1D, E). The increased puncta intensity was surprising because Laa1 localization is strongly reduced by the deletion of Laa2 ((Zysnarski et al., 2019) Fig. 2A,

**Figure 2.**
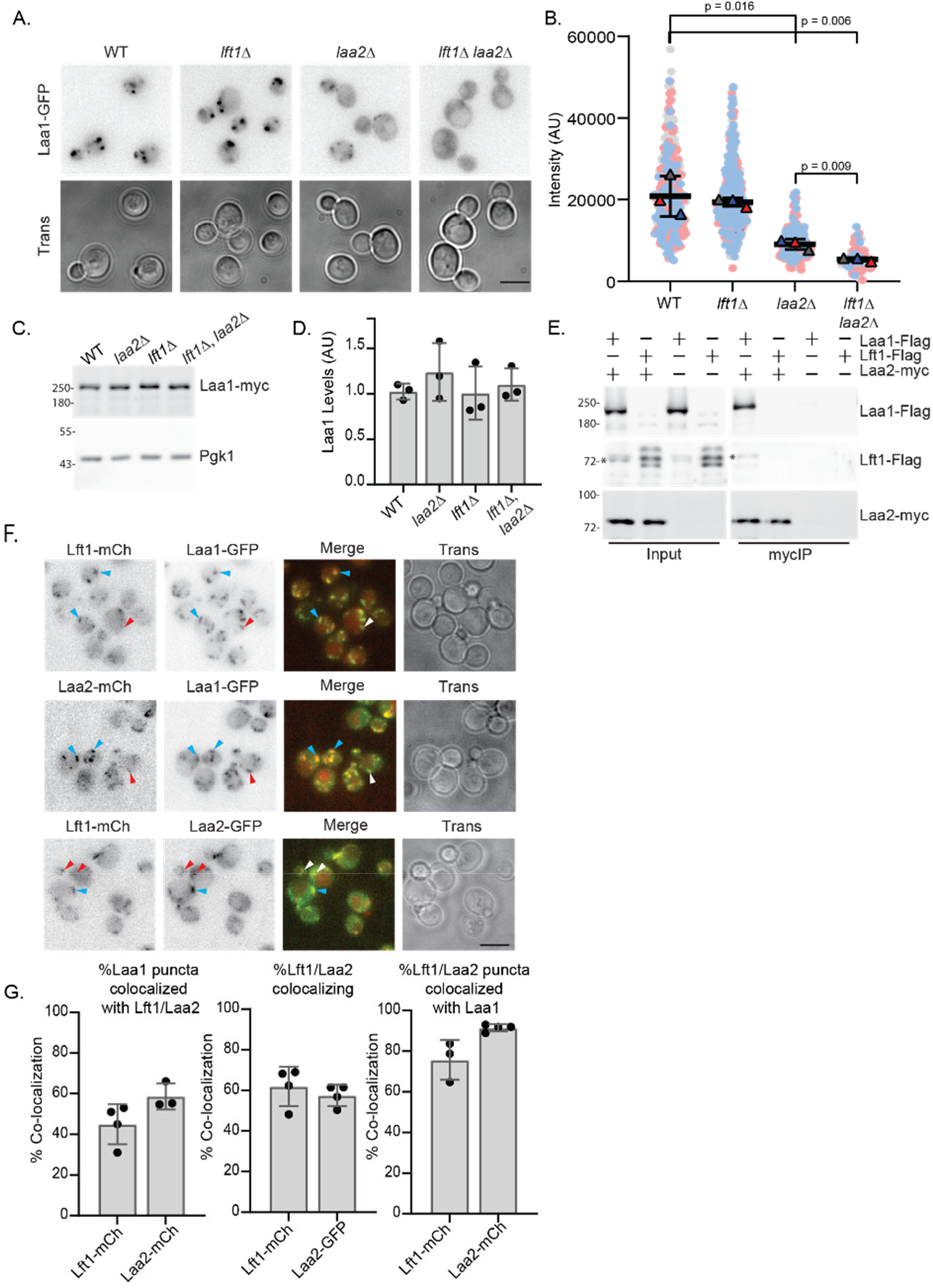
Laa1 forms mutually exclusive complexes with Lft1 and Laa2. A. Lft1 and Laa2 both contribute to Laa1 localization. Cells with indicated genotypes were imaged as described in materials and methods. B. Quantification of puncta intensity as described in Fig. 1C. C. Laa1 abundance is unaffected by loss of Lft1 or Laa2 alone or simultaneously. Lysates of cells of indicated genotypes were probed for myc or Pgk1, used as a lysis control. D. Quantification of data in C as described in Figure 1. E. Laa2 does not bind Lft1 as detected by immunoprecipitation. Myc-tagged proteins were immunoprecipitated from cells expressing indicated gene fusions from their endogenous loci and probed for myc and Flag. *indicates a degradation product from Laa1 migrating near the size of Lft1. Lft1-Flag migrates as a triplet under the conditions used for the myc IP F. Co-localization of Laa2, Laa1, and Lft1. Indicated gene fusions were expressed from their endogenous loci and cells were imaged as described in the materials and methods. Blue arrowheads indicate regions of co-localization. Red/White arrowheads indicate regions where co-localization was not observed. G. Percent of Laa1 structures that contained indicated protein (left). Percent of Lft1 structures that contained Laa2 (Lft1-mCh) or Laa2 structures that contained Lft1 (Laa2-GFP) (middle). Percent of Lft1 or Laa2 structures that contained Laa1 indicated protein that contained Charts show percentages from individual replicates, error bars indicate standard deviation. Micrographs show Z-stack compressions scale bars are 5 µm.

B). One possible explanation for this observation is that Lft1 and Laa2 interact with different pools of Laa1. To explore this possibility, we examined the localization of Laa1 in cells lacking Lft1 or Laa2 and in cells lacking both Lft1 and Laa2. We found that the loss of Lft1 alone did not cause a detectable effect on Laa1-GFP localization as monitored by puncta intensity (Fig. 2A, B). As previously reported, the loss of Laa2 reduced Laa1-GFP puncta intensity by over one-half ((Zysnarski et al., 2019), Fig. 2A, B). Notably, simultaneous loss of both Lft1 and Laa2 reduced Laa1-GFP puncta intensity to nearly undetectable levels (Fig. 2 A, B). This effect was not due to reduced Laa1 protein levels because Laa1 abundance was the same in wild-type cells and cells lacking Laa2 or Lft1 or both (Fig 2C, D). These findings suggest that Laa1 may exist in two populations, one bound to Lft1 and one bound to Laa2. Moreover, it indicates that Laa1 localization depends on these binding partners.

To determine whether two populations of Laa1 exist in cells, we asked whether Lft1 and Laa2 interact by co-immunoprecipitation. We were unable to detect an interaction between Laa2 and Lft1 under conditions where Laa2 binds Laa1(Fig. 2E). This indicates that Laa1 exists in two populations, one bound to Lft1 and one bound to Laa2.

To further test whether two populations of Laa1 exist in cells, we determined the extent to which Laa1 colocalized with Lft1 and Laa2. We found that approximately 50% of Laa1 structures colocalized with Laa2 and about 60% colocalized with Lft1, consistent with Laa1 existing in two populations. Similarly, approximately 60% of Lft1 structures contained Laa2 and vice versa. The partial co-localization is not due to the movement of the structures during image acquisition because Lft1 and Laa2 exhibit high levels of co-localization with Laa1. These data suggest that two populations of Laa1 exist in cells and that the two populations partially colocalize.

### Lft1 and Laa2 bind to an overlapping region on Laa1

To understand the structural basis of Laa1’s interaction with Lft1 and Laa2, we predicted complexes between Laa1 and Lft1 or Laa2 using AlphaFold2-based structure prediction. As illustrated in the PAE plots, which show the error of the prediction for the distance between every pair of residues in the structure (Sala et al., 2023; Varadi et al., 2022), Alphafold2 predicted both Lft1 and Laa2 contain small regions that make high-confidence contact with the C-terminus of Laa1 (Blue-Fig. 3A & B, Fig. S1 & 2). Surprisingly, despite little sequence similarity (Fig. 3C), the predicted interactions are highly similar (Fig. 3D). Both Lft1 and Laa2 contain several small alpha-helical regions separated by short, disordered linkers; these alpha-helical regions interact with an extended interface within the C-terminal 500 amino acids of Laa1 (Fig 3D). In the predictions, Lft1 binds to Laa1 via four alpha-helical regions (HR 1-4), whereas Laa2 binds to Laa1 via only three (HR1-3) (Fig 3C & D). Notably, in the predictions, HR1-3 of Lft1 and Laa2 superimpose almost exactly (Fig 3D). This result suggests that Lft1 and Laa2 bind to Laa1 via an overlapping interface and cannot simultaneously bind to Laa1.

**Figure 3.**
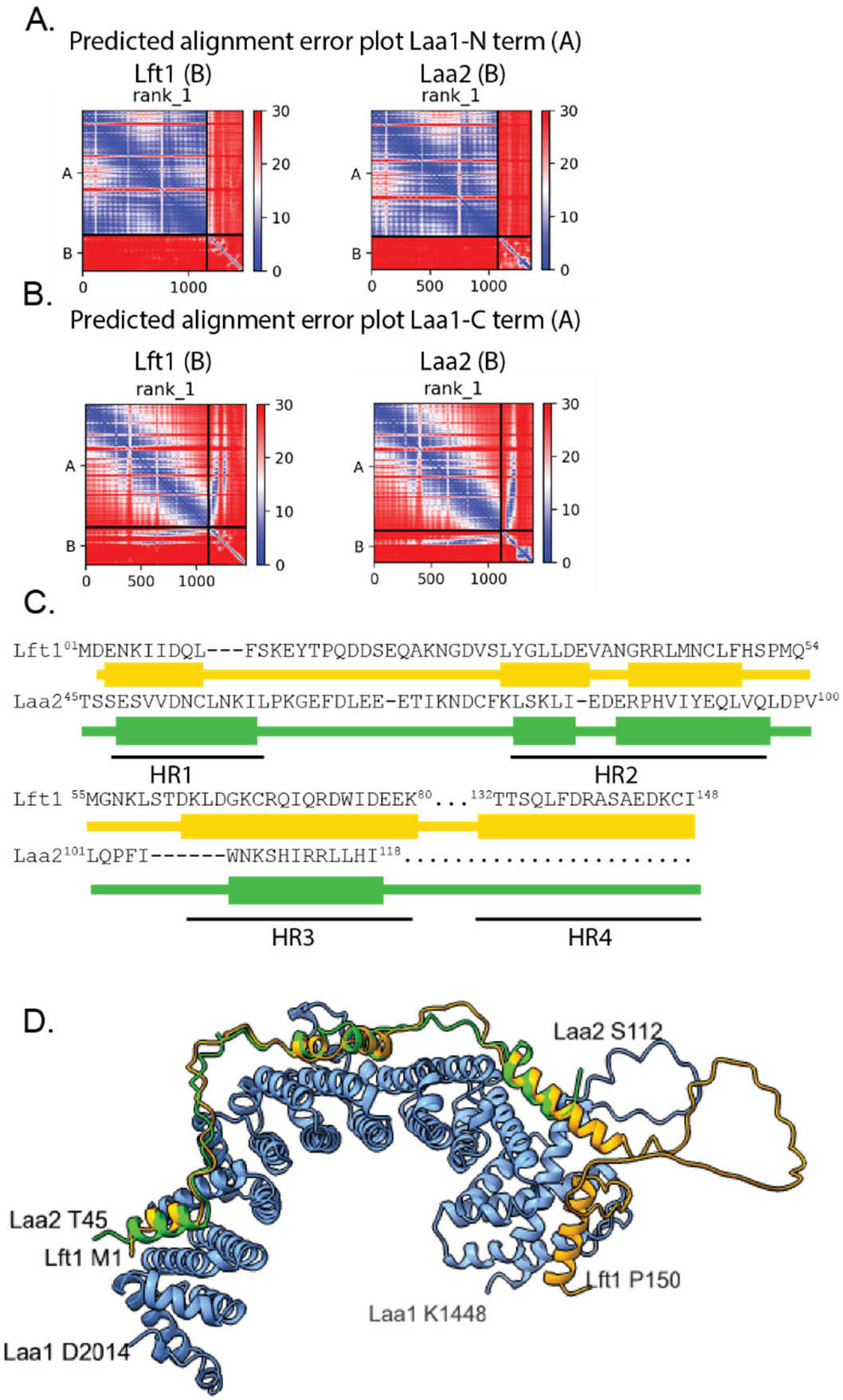
Lft1 and Laa2 are predicted to bind to an overlapping interface on the Laa1 C-terminus. A. No high- confidence interactions are predicted between the N-terminus of Laa1 and either Lft1 or Laa2, suggesting neither protein interacts with the N-terminus. Graphs show predicted alignment error for the top models of alpha fold prediction of dimer. Blue indicates the highest confidence. B. High confidence interactions are predicted between the C-terminus of Laa1 and both Lft1 and Laa2, suggesting both proteins interact with the C- terminus. Graphs as in A. C. Lft1 and Laa2 bear little sequence identity in predicted Laa1 interacting regions. The illustration shows sequence alignment of Lft1 and Laa2 based on overlay generated with ChimeraX matchmaker tool aligning the Laa1 C-terminus from the predicted dimeric structures. Helixes are indicated as boxes. The helical regions (HR) 1-4 are indicated below. D. Lft1 and Laa2 are predicted to adopt a similar fold when binding to Laa1. Illustration shows overlay of predicted structures described in C. Lft1 is orange, and Laa2 is green. Laa1 is in blue, the last 304 amino acids are indicated in a darker blue.

We validated the interaction interface of Laa1 with Lft1 and Laa2 using Laa1 truncations. We found that both Lft1 and Laa2 bound strongly to a fragment composed of the last 908 amino acids of Laa1 (pGal1- GST-Laa1(1106)), which contains the entire predicted binding domain (Fig. 4A-C). Conversely, neither protein bound to a Laa1 fragment that lacks the terminal 908 amino acids (Laa1(1106)-GST) (Fig. 4 D- F). These data indicate that C-terminal 908 amino acids of Laa1 are necessary and sufficient for binding to both Lft1 and Laa2. We next tested truncations that split the predicted binding interface. We found that a fragment composed of the last 304 amino acids of Laa1 (pGAL1-GST-Laa1(1710), which contains the binding interface for HR1 and truncates the binding interface for HR2, failed to bind either Lft1 or Laa2 (Fig. 4A-C). Finally, we found that a fragment of Laa1 that lacked the last 304 amino (Laa1(1710)-GST) bound weakly to Lft1, whereas it did not bind Laa2 (Fig. 4 D-F). Notably, this finding is consistent with the more extensive binding interface in the region N-terminal to amino acid 1710 predicted for Lft1 (HR3&4) compared to Laa2 (HR3). Finally, we note that Lft1 and Laa2 levels were lower in input samples expressing fragments of Laa1 lacking the binding sites, consistent with the observation that Lft1 and Laa2 are destabilized in the absence of Laa1 (Fig 1 D (Zysnarski et al., 2019)). Conversely, Lft1 levels were elevated in samples overexpressing Laa1 fragments that contained the full binding site (Fig. 4B). The concordance of our experimental data with the predicted structure strongly suggests the prediction captures the binding interface on Laa1 accurately. Together, these results suggest that Lft1 and Laa2 cannot bind Laa1 simultaneously and indicate that Lft1 defines a second Laa1-containing complex in budding yeast.

**Figure 4.**
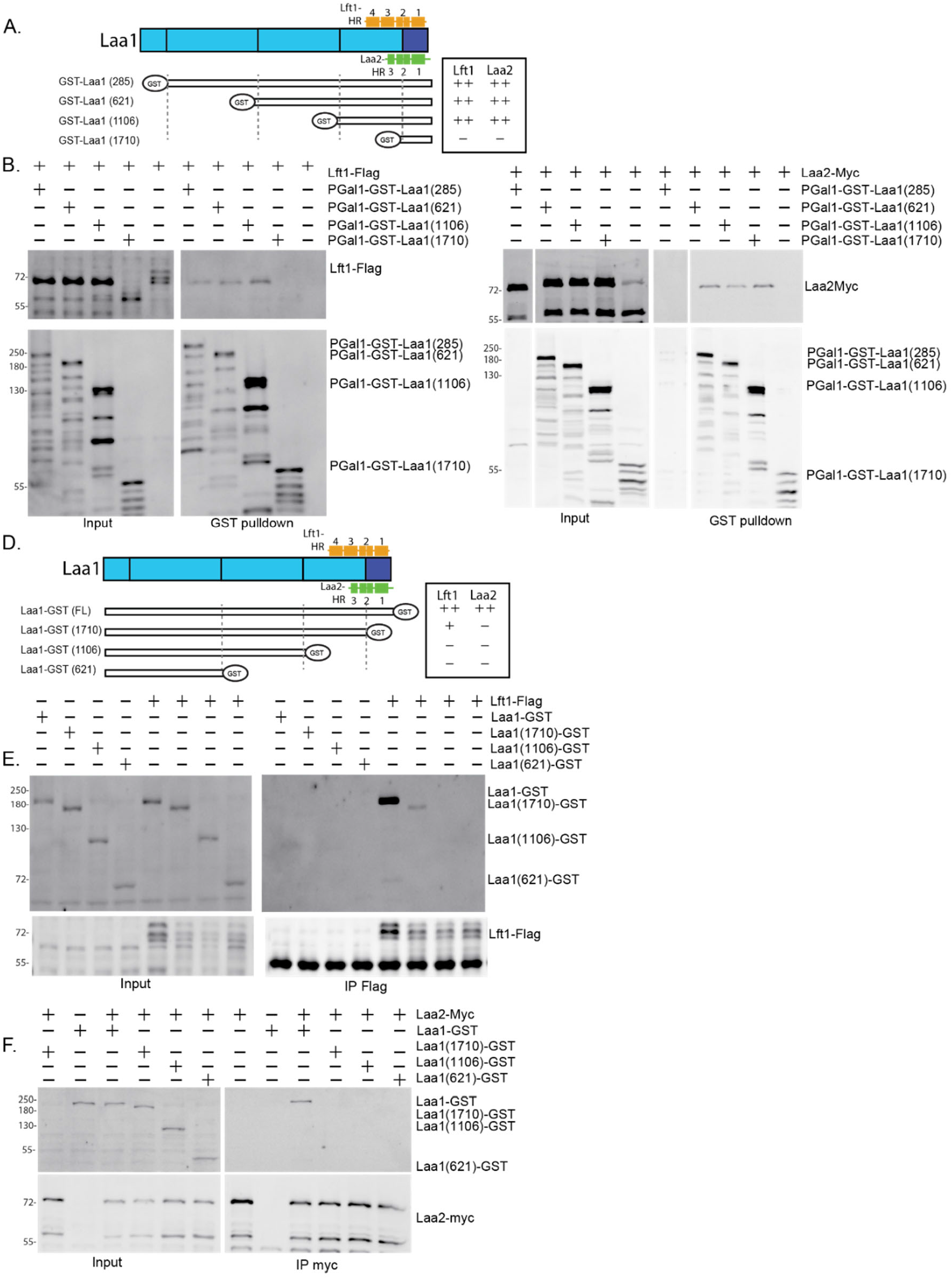
Lft1 and Laa2 bind to the C-terminus of Laa1. A. Schematic of Laa1 fragments used in B and C, and approximate orientation to Lft1 and Laa2 HRs. Strength of binding to Lft1 and Laa2 is indicated to the right. B & C Lft1 and Laa2 bind to Laa1 fragments that contain the last 908 amino acids of Laa1 (pGAL1-GST-Laa1 (1106). GST-tagged proteins were isolated using Glutathione beads from cells expressing indicated gene fusions expressed from their endogenous loci and probed for GST and Flag or Myc. Note that Lft1-Flag migrates as a single band under conditions where Laa1 is overexpressed and can bind Lft1, it migrates as a triplet when Laa1 is not overexpressed. D. Schematic of Laa1 fragments used in E and F and approximate orientation to Lft1 and Laa2 HRs. Strength of binding to Lft1 and Laa2 is indicated to the right. E. Lft1 binds weakly to Laa1 missing the last 304 amino acids (Laa1(1710)-GST), but not larger deletions. Flag-tagged proteins were immunoprecipitated from cells expressing indicated gene fusions expressed from their endogenous loci and probed for GST and Flag. F. Laa2 does not bind to any C-terminal truncations tested. Myc-tagged proteins were immunoprecipitated from cells expressing indicated gene fusions expressed from their endogenous loci and probed for GST and myc.

To validate the predicted interface on Lft1, we generated mutations in Lft1 that removed HR1, HR2, and HR3 individually. We found that removing either HR1 or HR3 in Lft1 expressed from its endogenous locus reduced the interaction of Lft1 with Laa1 to about 60% of wild-type levels (Fig. 5A, B). However, removing HR2 reduced the interaction to undetectable levels (Fig. 5A, B). Together these results support the prediction that the Lft1 binding interface is extended and utilizes contacts spanning the region containing HR1-4.

**Figure 5.**
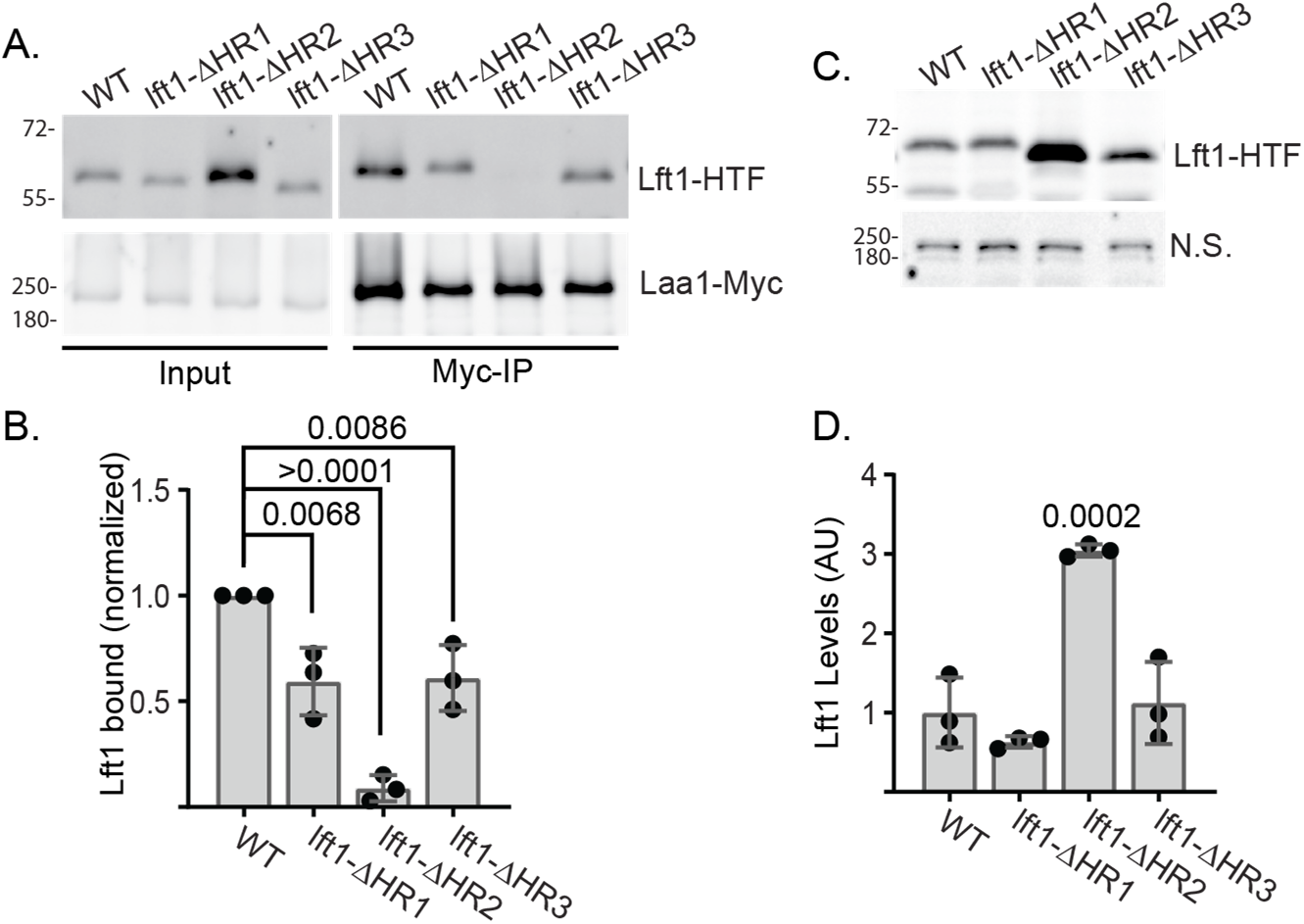
Multiple Lft1 helical regions contribute to binding. A. Indicated *LFT1* mutations were tagged with 6-HIS-TEV-3FLAG (HTF) and expressed from the endogenous locus in cells expressing Laa1-myc from the endogenous locus. Myc-tagged proteins were immunoprecipitated and probed for Myc and Flag B. Lft1 bound was quantified. Lft1 signal was normalized to the amount of Laa1 in each sample and then to the normalized signal for wild-type reactions on the same immunoblot. C. lft1-ΔHR2 is more abundant than wild-type Lft1. Lysates from strains described in A were generated and immunoblotted for Flag. Equal loading was confirmed with an endogenous protein that cross-reacts with the flag antibody (N.S.) D. Lft1 abundance was quantified. Signal intensity was normalized to the average of the wild-type signal. P-values derive from Dunnett’s multiple comparisons test. Charts show standard deviations.

Curiously, Lft1-ΔHR2 is 3-fold more abundant than wild-type Lft1, whereas the abundance of Lft1-ΔHR1 and Lft1-ΔHR3 is not significantly different from that of wild-type Lft1 (Fig. 5C, D). This finding is surprising because Lft1 is less abundant in cells lacking Laa1 or in cells expressing truncations of Laa1 that do not bind Lft1, suggesting that Lft1 is destabilized when it does not bind Laa1. The increased abundance of Lft1-ΔHR2 suggests that HR2 may play a role in destabilizing Lft1 when it is not bound to Laa1, potentially by encoding a degron or interaction interface for proteins that target it for degradation.

### The Lft1-Laa1 complex does not bind AP1 or control its localization

The only molecular function known for Laa1 is to recruit AP1, a function described for HEATR5 proteins in many other species (Gillard et al., 2015; Le Bras et al., 2012; Madan et al., 2023; Yu et al., 2012; Zysnarski et al., 2019). However, HEATR5-proteins are not thought to bind directly to AP1. Instead, they rely on accessory proteins that contain AP1 binding sites that mediate the interaction between the HEATR5 protein and AP1. The AP1 binding site often takes the form of a small peptide motif with the consensus FXXφ (where F is phenylalanine, X is any amino acid, and φ is a hydrophobic residue) in a region rich in acidic amino acids (Duncan et al., 2003; Duncan and Payne, 2003; Mattera et al., 2004). We previously reported that a mutation of the FXXφ in Laa2 eliminated the interaction between Laa1 and AP1 (Zysnarski et al., 2019). However, Lft1 contains one potential AP1 binding motif in its unstructured C-terminus (^298^DDGDGFEQV), raising the possibility that it could bind AP1. Therefore, we tested whether Lft1 binds AP1. Surprisingly, we were unable to detect an interaction between Lft1 and AP1 using 2-hybrid analysis or co-immunoprecipitation under conditions where Laa2 binds AP1 (Fig. 6 A-C). These data suggest that if Lft1 interacts with AP1, the interaction is weak or depends on other factors that are not recapitulated in our assays.

**Figure 6.**
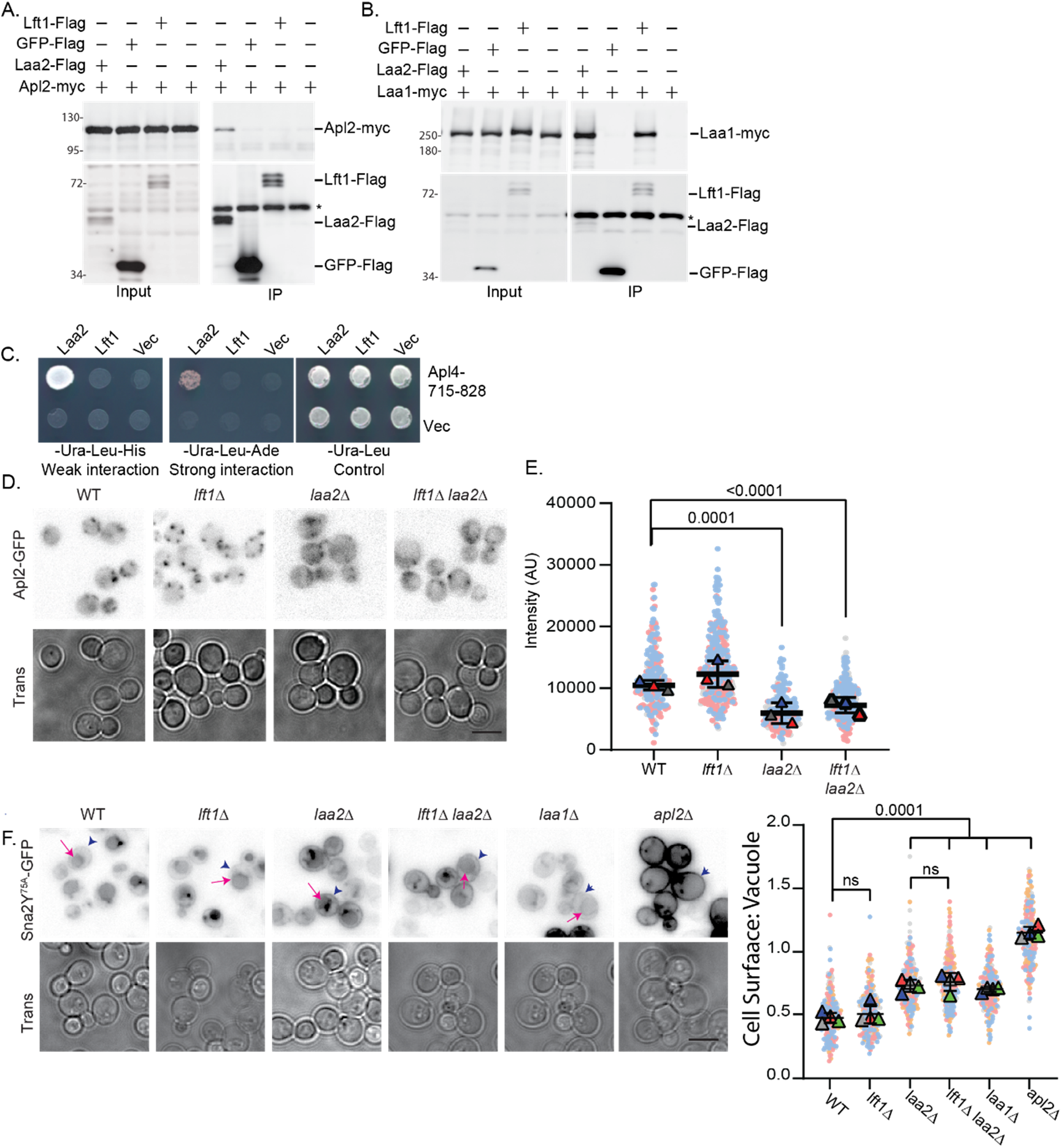
Lft1 does not detectably bind to AP1 or contribute to its localization or function in sorting Sna2^Y75A^. A. AP1 was not detected in coimmunoprecipitation samples with Lft1 under conditions where Laa2 binds AP1 and B. under conditions where Lft1 binds Laa1. Myc-tagged proteins were immunoprecipitated from cells expressing indicated gene fusions expressed from their endogenous loci and probed for Myc and Flag. *indicates antibody band. C. An interaction between Lft1 and AP1-γ ear (Apl4-a.a.715-828) was not detected by 2-hybrid under conditions where AP1- γ ear bind Laa2. D. Loss of Lft1 does not impact AP1 localization. Cells with indicated genotypes were imaged. E. Quantification of puncta intensity as described in Fig. 1C. F. Lft1 does not contribute to Sna2^Y75A^ sorting. Cells with indicated genotypes expressing GFP-Sna2^Y75A^ from a plasmid were imaged. Blue arrowheads indicate plasma membrane localization and magenta arrows indicate plasma membrane localization. G. Quantification of the plasma membrane to vacuole intensity for single cells. P-values derive from Dunnett’s multiple comparisons test. Chart shows standard deviations of the mean for replicate experiments. Scale bars are 5 µm.

To investigate whether the Lft1-Laa1 complex participates in AP1 recruitment, we examined AP1 localization in cells lacking Lft1. We found that the loss of Lft1 did not affect AP1 localization as monitored by puncta intensity, and Lft1 did not enhance the AP1 localization defect caused by the loss of Laa2 (Fig. 6 D, E). These results argue that, unlike the Laa2-Laa1 complex, the Lft1-Laa1 complex is not important for recruiting or stabilizing AP1 on membranes. We next asked whether the Lft1-Laa1 complex depends on AP1 for its localization. We found that loss of the AP1 subunit Apl4 elevated Lft1 puncta intensity, similar to the effect of Laa2 (Fig. 1B). The similar effect of *apl2Δ* and *laa2Δ* may indicate that the elevated intensity of Lft1 puncta is due to loss of AP1 function, for example by delaying TGN maturation, rather than competition between Laa2 and Lft1. In any case, these results argue that AP1 does not recruit the Lft1-Laa1 complex to membranes. These findings suggest that Lft1-Laa1 performs at least one function besides recruiting AP1 to membranes.

As a further assessment of the role of Lft1 in AP1-mediated traffic, we examined the effect of Lft1 on Sna2^Y75A^. Sna2 is a vacuolar membrane protein that normally localizes to the vacuole. Its delivery to the vacuole depends on two tyrosine motifs Y75A, which is recognized by AP3, and Y65A, which is recognized by the AP1-μ-subunit encoded by *APM1* (Renard et al., 2010; Whitfield et al., 2016). Defects in AP1 cause a complete redistribution of Sna2^Y75A^-GFP to the plasma membrane, whereas deletion of *laa1Δ* or *laa2Δ* cause a partial relocalization of Sna2^Y75A^-GFP to the plasma membrane, likely reflecting the strong but not complete mislocalization of AP1 in *laa1Δ* or *laa2Δ* cells (Zysnarski et al., 2019). We assessed the localization of Sna2^Y75A^ expressed from a plasmid (Fig. 6F & G). To control for different levels of expression based on plasmid copy number, we compared the intensity of the plasma membrane signal to the intensity of the vacuolar signal in the same cell. Based on this analysis, we found that *lft1Δ* did not elevate Sna2^Y75A^ levels at the plasma membrane, even when combined with *laa2Δ* (Fig. 6G). These results suggest that Lft1 is not involved in the traffic of Sna2^Y75A^, a direct cargo of AP1 (Whitfield, 2018; Whitfield et al., 2016). These data indicate that the Lft1-Laa2 complex performs a different function than the Laa2-Laa1 complex.

### The Lft1 complex localizes to the late TGN

The TGN in yeast is a dynamically maturing compartment. Its maturation state is associated with waves of traffic, which are mediated by different coats and responsible for cargo exit from or retention in the TGN (Casler et al., 2019; Daboussi et al., 2012; Highland and Fromme, 2021; Tojima et al., 2019). Lft1 is highly dynamic and expressed at relatively low levels. For this reason, it was difficult to track it accurately in time-lapse microscopy. Therefore, we turned to co-localization analysis to determine whether Lft1 is associated with TGN at a specific time during their maturation. When coupled with fiduciary markers that are known to be enriched at different stages of maturation, this is an effective method for positioning proteins to stages within maturing compartments that has been applied to Golgi and endosome maturation (Arlt et al., 2015; Kim et al., 2016). The maturation stage of the TGN is marked by different clathrin adaptors. The clathrin adaptor Gga2 is enriched on early TGN, whereas the clathrin adaptors Ent5 and AP1 are more enriched on more mature TGN (Casler et al., 2019). To determine whether Lft1 is associated with nascent or more mature TGN, we monitored the co-localization of Lft1 with the different adaptors. We found that Lft1 colocalized better with Ent5 and AP1 than with Gga2 (Fig. 7 A & B). These findings suggest that Lft1 is enriched on TGN at later stages of maturation.

**Figure 7.**
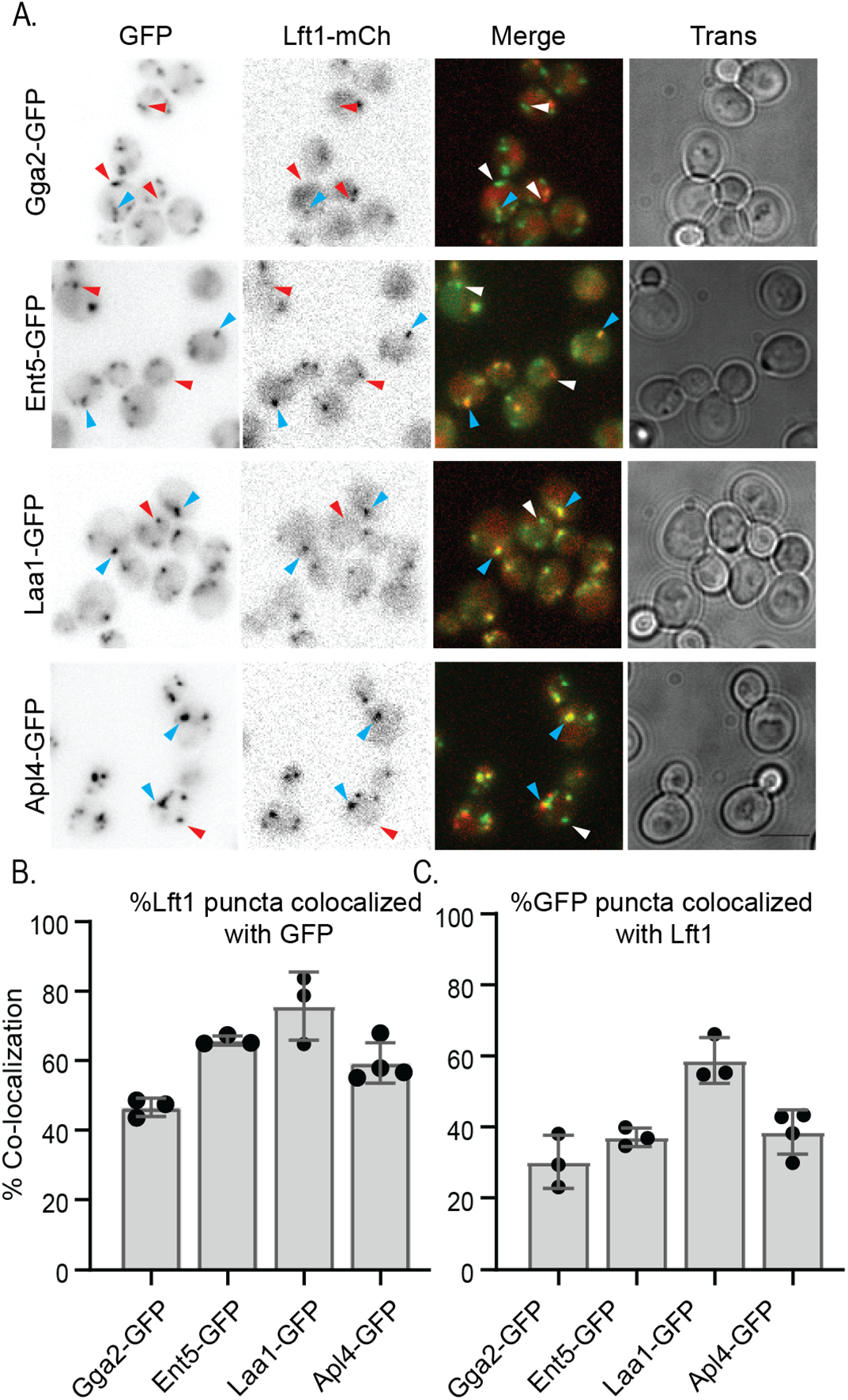
Lft1 shows a higher level of co-localization with late stage TGN. A. Indicated gene fusions were expressed from their endogenous loci and cells were imaged as described in the materials and methods. Scale bar is 5 μm. Blue arrowheads indicate regions of co-localization. Red/White arrowheads indicate regions where co-localization was not observed. B. Percent of Lft1 structures that contained indicated GFP protein and C. Percent of structures labeled with indicated GFP-protein that contained Lft1 were determined as described in materials and methods. Charts show percentages from individual replicates, error bars indicate standard deviation.

### The Lft1 complex mediates recycling in the TGN

To better understand the function of the Laa1-Lft1 complex, we examined the effect of the loss of Lft1 alone or in combination with Laa2 on phenotypes associated with the loss of Laa1. We first examined Sertraline sensitivity. Sertraline is a cationic amphiphilic drug that causes severe growth defects when combined with mutations that impair traffic at the TGN or in the early endosomal system (Rainey et al., 2010). We previously determined that cells lacking Laa1 or Laa2 are sensitive to Sertraline (Zysnarski et al., 2019). However, cells lacking Laa2 were slightly less sensitive than those lacking Laa1.

Moreover, in a high throughput study, cells lacking Laa1, Laa2, or Lft1 were all more sensitive to Sertraline than wild-type cells. However, cells lacking Laa1 were far more sensitive than those lacking either Laa2 or Lft1. When ranked in order of the growth defect caused by 12.5 µM Sertraline *laa1Δ* was ranked 16^th^ out of 4198, whereas *laa2Δ* was the 54^th^ most sensitive, and *lft1Δ* was the 606^th^ most sensitive to Sertraline (Ericson et al., 2008). These differences in sensitivity suggest the Laa2-Laa1 and Lft1-Laa1 complex might both contribute to the Sertraline sensitive function of Laa1. To confirm this, we examined the Sertraline sensitivity of cells lacking Laa1, Lft1, Laa2, or a combination of Lft1 and Laa2. Consistent with the prior results, the *laa1Δ* cells were more sensitive than the single deletion of either *LFT1* or *LAA2* as monitored by the IC50 of Sertraline for each of the mutant genotypes (Fig. 8A). Importantly, consistent with the hypothesis that both the Lft1-Laa1 and Laa2-Laa1 complexes contribute to Sertraline sensitivity, the double *lft1Δ laa2Δ* genotype was nearly as sensitive to Sertraline as the *laa1Δ* genotype (Fig. 8A).

**Figure 8.**
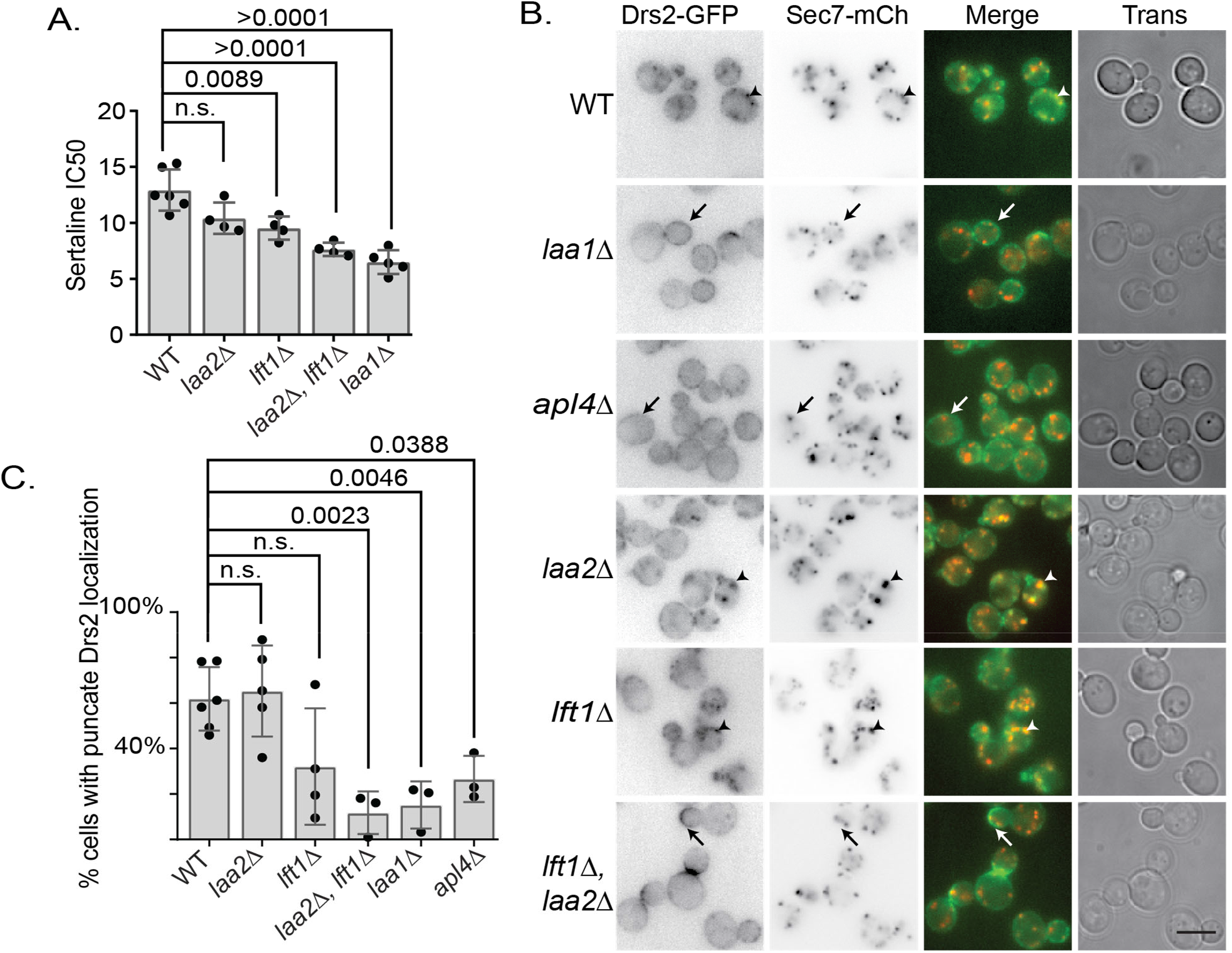
The Lft1-Laa1 and Laa2-Lft1 complexes both mediate intra-Golgi recycling. A. Cells lacking both Lft1 and Laa2 are sensitive to Sertraline at levels similar to cells lacking Laa1. Charts show IC50 values from replicate experiments and standard deviations. P-values are from a student t-test. B. Cells lacking both Lft1 and Laa2 are severely defective in Drs2 recycling. Drs2 recycling was monitored. Cells with defective recycling accumulate Drs2 at the cell surface after latrunculin A treatment (arrows), whereas those with normal recycling have Drs2 in Sec7-positive puncta (arrowheads) C. Quantification cells with punctate Drs2 localization as described in the materials and methods. Charts show the percent of cells scored as having defects in Drs2 localization from replicate experiments and standard deviations. P-values are from a student t-test. Scale bar is 5 μm.

The growth defect of the double *lft1Δ laa2Δ* mutation in Sertraline suggests that both complexes contribute to Laa1 functions. However, although Sertraline sensitivity is a quantitative metric of defects, the mechanism of Sertraline sensitivity is unclear. Therefore, we examined the effect of Lft1 and Laa2 in traffic at the TGN. To do this, we monitored the localization of Drs2, an aminophospholipid translocase that localizes to the TGN (Chen et al., 1999). The TGN in yeast is a maturing compartment. Therefore, the steady-state localization of TGN-resident proteins like Drs2 requires their retrieval from late-stage compartments to early-stage compartments. This ‘intra-Golgi’ recycling mechanism depends on the clathrin adaptors AP1 and Ent5 (Casler et al., 2022). When intra-Golgi recycling is impaired, Drs2 is transported to the plasma membrane. It is then rapidly endocytosed and sorted to the TGN (Liu et al., 2008). Therefore, to visualize defects in Drs2 intra-Golgi recycling, we inhibited endocytosis by treating cells with Latrunculin A as previously described (Liu et al., 2008). Under these conditions, in wild-type cells, a substantial fraction of Drs2 remained localized to the TGN as assessed by co-localization with Sec7 (Fig. 8B). In contrast, in cells lacking either AP1 or Laa1, most of the Drs2 is at the cell surface under these conditions (Fig. 8B).

To quantify the phenotype, we performed a blinded analysis of whether Drs2 localized to puncta or the plasma membrane in the different samples. In wild-type cells treated with Latrunculin A, approximately 60% of the cells were classified as punctate using this analysis. In contrast, in *apl4Δ* or *laa1Δ* cells treated with Latrunculin A, this number was 20% and 15%, respectively (Fig. 8C). Loss of Laa2 alone had little effect on the percent of cells with punctate Drs2 in cells treated with Latrunculin A. In contrast, loss of Lft1 caused a variable outcome. However, in cells lacking both Ltf1 and Laa2, the percent of cells with punctate localization was significantly lower than wild-type and reached a level similar to *laa1Δ* cells. These data suggest that both the Lft1-Laa1 and the Laa2-Laa1 complexes participate in intra-Golgi recycling to maintain Drs2 at the TGN, even though the Lft1-Laa1 complex does not impact AP1 localization.

To determine whether the effect on Drs2 was due to a general defect of the TGN we examined the effect of the loss of Laa1, Laa2, and Lft1 on AP3-mediated traffic. To do this, we monitored the vacuolar localization of GFP-NSI, a synthetic cargo whose localization to the vacuole depends on the AP3 pathway (Plemel et al., 2021). In cells lacking AP3 (*apl6Δ*), GFP-NSI is missorted to the plasma membrane (Fig. S3). In contrast, in wild-type cells and cells lacking Laa1, Laa2, Lft1, or AP1 (*apl4Δ*), GFP-NSI is delivered to the vacuole. These data suggest that neither Laa1 complexes contribute to AP3-mediated traffic and that their loss does not broadly disrupt traffic at the TGN.

Because Ent5 is also important for Drs2 localization, we tested if Lft1 interacted with Ent5 (Casler et al., 2022). We failed to detect an interaction of Lft1, Laa1, or Laa2 with Ent5 under conditions where Ent5 binds to AP1 (Fig. S4). These data suggest that if either Laa1 complex interacts with Ent5 the interaction is weak or depends on other factors that are not retained during immunoprecipitation.

### Human proteins are predicted to bind to Heatr5b via a conserved mechanism

Although HEATR5 proteins themselves are slowly evolving, their binding partners are fast-evolving (Kuznetsov et al., 2023). For example, clear homologs of Lft1 and Laa2 can be identified only in other fungi (Huerta-Cepas et al., 2019). Similarly, clear homologs to the human Heatr5b binding proteins Aftiphilin and γ-synergin can be identified only in animals (Huerta-Cepas et al., 2019). Since Lft1 and Laa2 bear little sequence similarity yet bind Laa1 in a similar manner, we asked whether other known or suspected HEATR5 binding proteins might share a binding mechanism. To do this, we used AlphaFold2 to predict interactions between Heatr5b and its known binding proteins Aftiphilin and γ-synergin, as well as Clba1, an uncharacterized protein with weak similarity to a region of Aftiphilin and Laa2 (Zysnarski et al., 2019). We also predicted interactions between Heatr5a, a paralog of Heatr5b of unknown function that is only 58% identical to HEATR5b. Alphafold2 did not yield high-confidence interactions between Heatr5a or Heatr5b and γ-synergin (unpublished data). However, Alphafold2 predicted high-confidence interactions between the C-termini of Heatr5a and Heatr5b and both Aftiphilin and Clba1 (Fig. 9A, Fig. S5-S8). Aftiphilin and Clba1 are predicted to bind over an extended interface in the C-termini of Heatr5a and Heatr5b via three short alpha-helical regions analogous to HR1-3 in Lft1/Laa2. These regions in Aftiphilin and Clba1 have weak but identifiable sequence similarities to one another and to Laa2 (Fig. 9B, (Zysnarski et al., 2019)). Indeed, an area overlapping with HR2 and HR3 has been annotated as the Clba (clathrin binding of Aftiphilin) domain for which Clba1 was named by the HUGO Gene Nomenclature Committee at the European Bioinformatics Institute (Thomas et al., 2022). As with the budding yeast proteins, the human proteins superimpose almost exactly with one another, suggesting that they cannot bind to HEATR5 proteins simultaneously (Fig. 9c). We also compared the predicted interaction surfaces between the budding yeast and human complexes. When aligning the predicted complexes based on the similarity of Laa1 and HEATR5b, we found that Lft1 and Aftiphilin superimposed well, particularly in HR2 and HR3 (Fig. 9d). Together, these results suggest that multiple HEATR5 binding proteins compete for the same interface in both budding yeast and humans and suggest that human HEATR5 proteins may form two or more biochemically distinct complexes.

**Figure 9.**
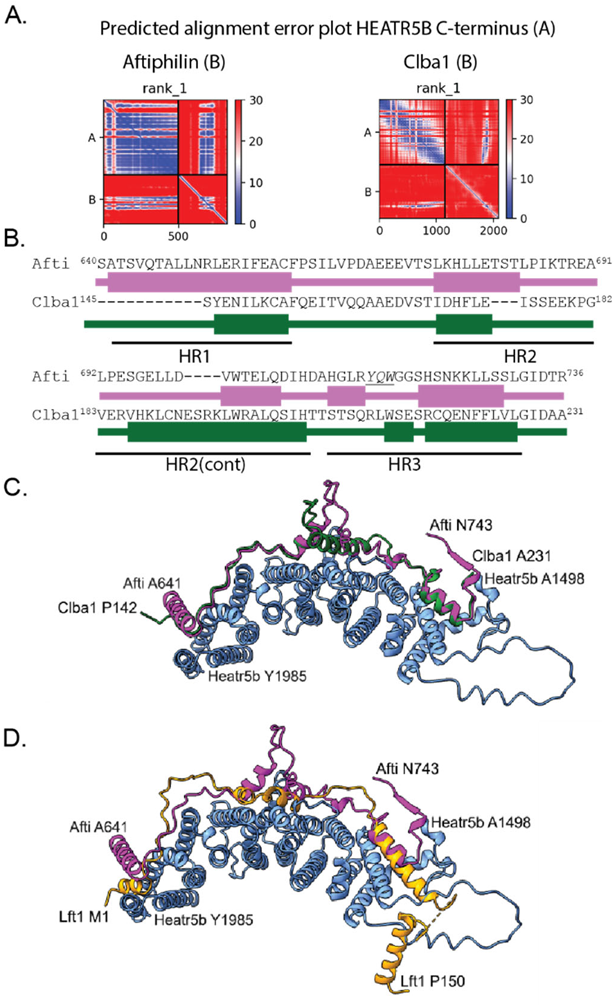
Human proteins are predicted to bind an overlapping interface in the c-terminus of Heatr5b similar to the Laa1-Lft1/Laa2 binding interface. A. High confidence interactions are predicted between the C- terminus of Heatr5b and Aftiphilin and Clba1. Charts are as described in Figure 4. B. Aftifphilin and Clba1 display weak similarity in the predicted Heatr5 interacting regions. Diagrams are as described in Figure 4, based on matchmaker tool aligning Aftiphilin and Clba1 in the predicted structures. C. Aftilphilin and Clba1 are predicted to adopt a similar fold and interact with the same regions of Heatr5b. Illustration shows structure alignment described in panel B. D. Aftiphilin and Lft1 are predicted to bind HEATR5 proteins via a related interface. Illustration shows structure alignment based on matchmaker tool aligning Laa1 and HEATR5b. Laa1/HEATR5b are in blue, Aftiphilin is in magenta, Clba1 is in dark green and Lft1 is in orange.

## Discussion

In this study, we report the discovery that the yeast HEATR5 protein, Laa1, participates in two distinct biochemical protein complexes. This is the first report of any HEATR5 protein participating in distinct biochemical complexes with mutually exclusive binding partners. Our discovery challenges previous models for Laa1 functions and introduces the concept that HEATR5 proteins may participate in multiple distinct complexes in other organisms.

Our findings indicate that the Lft1-Laa1 and Laa2-Laa1 complexes are at least partially redundant. Multiple models could explain this redundancy. The two complexes may function interchangeably at the same step (Fig. 10A). Alternately, the two complexes could act in different recycling pathways that emerge from different domains of the TGN or different stages of TGN maturation (Fig. 10B & C). Future work is needed to distinguish these possibilities.

**Figure 10.**
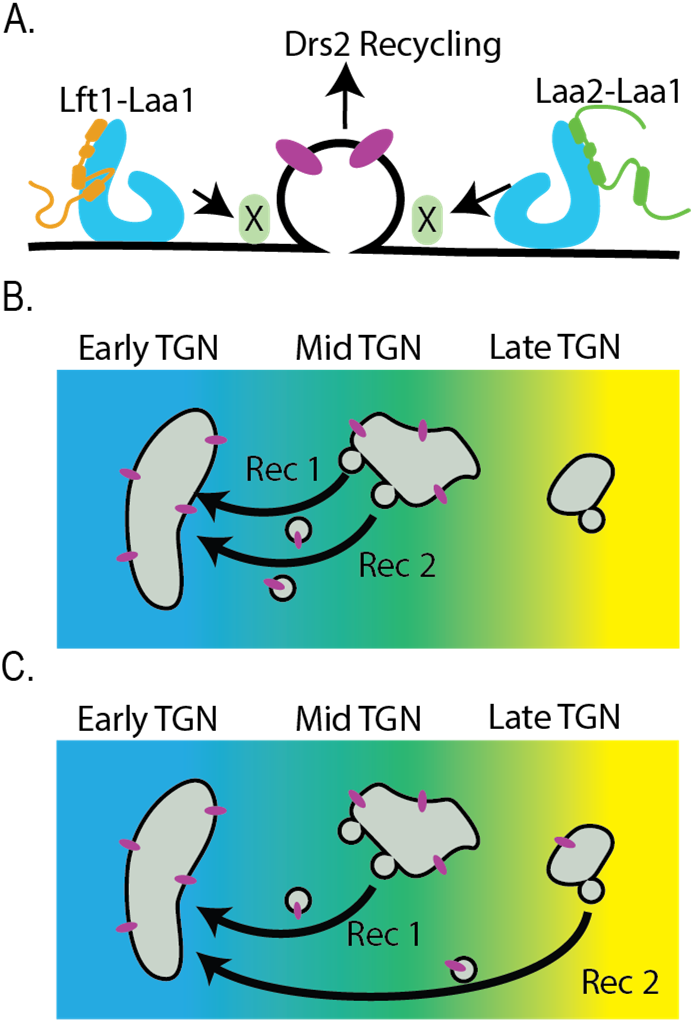
Models of potential roles of Lft1-Laa1 and Laa2-Laa1 complexes in intra-Golgi recycling. A. The two complexes redundantly perform an activity needed to recycle Drs2 from a late-stage TGN to an early-stage TGN. B and C. The two complexes act in two different pathways (Rec 1 and Rec 2) that emerge from different domains of the TGN (B) or that act on sequential stages of TGN maturation (C). Drs2 can recycle by either pathway. When both are disrupted Drs2 is transported to the cell surface.

Considering our findings that Laa1 performs a function besides recruiting AP1, a re-reading of the prior literature, particularly analysis of high-throughput studies, reveals that other HEATR5 proteins may also have functions besides recruiting AP1. In human cells, knocking down Heatr5b or Aftiphilin causes endocytosed transferrin to accumulate in peripheral endosomes, a phenotype not observed with AP1 knockdown (Hirst et al., 2005). In worms, the HEATR5 homolog SOAP1, but not AP1, was identified in a screen for genes important for endocytosis (Balklava et al., 2007). In addition, the obligate Heatr5b interaction partners Aftiphilin and γ-synergin were found in the interactome of AP2 and AP3 in human cells, and the plant HEATR5 protein, Sweetie, was found in the interactome of the endocytic T-plate complex (Burman et al., 2005; Dragwidge et al., 2023; Stockhammer et al., 2022). These studies suggest HEATR5 proteins in other organisms have roles outside the AP1 pathway. Whether these additional roles involve two or more distinct HEATR5 complexes remains to be determined.

This study also defines the common molecular binding interface between the yeast HEATR5 protein and its co-factors using high-confidence structure predictions and biochemical analyses. In addition, we report high-confidence predictions of Heatr5a and Heatr5b complexes with Aftiphilin and Clab1. We find that the predicted human complexes harbor interaction interfaces highly similar to the experimentally verified interface in the yeast proteins.

Notably, a published dissertation supports these predictions (Whitfield, 2018). It reports that Clba1 binds Heatr5b when overexpressed in HEK293T cells. Moreover, the work identifies two additional proteins, Fez1 and Fez2, that contain limited homology to Laa2, including the region we identify as mediating Laa1 binding. Fez1 and Fez2 are ∼50% identical. Both Fez1 and Fex2 bind Heatr5a when overexpressed. Importantly, mutations that lie between the regions we refer to as HR2 and HR3 disrupt the interactions of Aftiphilin and Clba1, and Fez2 with Heatr5b or Heatr5a respectively. These data indicate that Heatr5 proteins in both yeast and humans use a similar binding interface for their co-factors. Furthermore, these data suggest that both Heatr5a and Heatr5b bind mutually exclusive co-factors and exist in multiple biochemically distinct subcomplexes. Significantly, flies and worms contain one HEATR5 protein but contain homologs of both Fez1 and Aftiphilin. This suggests that these other HEART5 proteins also function in multiple distinct subcomplexes.

The interaction between Laa1 and Lft1 or Laa2 is stable enough to be maintained during immunoprecipitations (Fig. 1, (Zysnarski et al., 2019)). However, it remains unclear how stable the interactions are in vivo. Laa1 binding protects both wild-type Lft1 and Laa2 from degradation, as seen by the lower abundances of both in cells lacking Laa1 (Fig. 1, (Zysnarski et al., 2019)). This lower abundance argues that if Lft1-Laa1 and Laa2-Laa1 complexes dynamically assemble and disassemble, free Lft1 and Laa2 are likely short-lived either because the released protein is rapidly bound to a new Laa1 molecule or degraded. Notably, Lft1 and Laa2 have different degradation kinetics. In a genome-wide analysis of protein stability, Laa2 has a half-life of 10.1 hours, similar to the half-life of Laa1 (10.5 hr) and the bulk of the budding yeast proteome (>8 hr), whose abundance is regulated by synthesis rate rather than degradation rate (Christiano et al., 2014). This half-life is consistent with Laa2 either binding stably to Laa1 or rapidly rebinding such that Laa2 is not actively degraded even if it engages in dynamic assembly cycles. In contrast, Lft1 has a half-life of 5.7 hours, placing it in the 15% budding yeast proteome specifically targeted by degradation pathways (Christiano et al., 2014). Future work will be needed to assess whether Lft1-Laa1 or Laa2-Laa1 complexes assemble and disassemble dynamically in vivo and, if so, whether targeted degradation of Lft1 plays a vital role under normal conditions.

The extended nature of the Laa1 binding site on Lft1 and Laa2 may be significant for their molecular mechanism. The Lft1 and Laa2 binding sites for Laa1 are composed of a series of alpha-helical regions separated by unstructured segments (Fig. 3). Our studies using truncations of Laa1 and HR deletion mutations of Lft1 indicate that multiple areas of this extended interface contribute to the binding mechanism (Fig. 4, 6). We speculate that this interface may allow Lft1 and Laa2 to remain bound to Laa1 even if the conformation of Laa1 changes. Based on its predicted structure, Laa1 is likely to be conformationally dynamic. The predicted C-terminus of Laa1 is almost entirely tandem HEAT repeats. The HEAT repeat is a simple fold composed of two alpha helices separated by a short linker. This arrangement is common in biology. Tandem HEAT repeats appear in diverse proteins, including kinases, importins, transcription factors, and vesicular coat proteins (Kobe et al., 1999). In these proteins, the tandem HEAT repeats form an alpha-solenoid that can display an extraordinary level of conformational flexibility, in some cases elastically stretching by 2-fold (Forwood et al., 2010; Kappel et al., 2010; Yoshimura and Hirano, 2016). In addition, Laa1 contains an unstructured loop between amino acids 1711- 1733, which lies near the binding sites for HR2 on Lft1 and Laa2. If Laa1 undergoes elastic stretching or other conformational changes, the extended binding interface may play a role in maintaining the interaction between Laa1 and its binding partners.

In summary, these findings reveal that Laa1 participates in two biochemically distinct complexes. The complexes are defined by the mutually exclusive binding of two fast-evolving binding partners. The binding mechanism is characterized by an extended surface on both Laa1 and its binding partners. The binding mechanism may provide a paradigm for HEATR5 binding proteins in many organisms. The two Laa1 complexes function at least partially redundantly in intra-Golgi recycling, however, their exact mechanistic function remains unknown. Given the large size of Laa1, there are likely additional protein- protein or protein-lipid interactions mediated by Laa1 that remain undiscovered. Future studies are needed to elucidate these additional interactions and reveal a complete picture of the molecular mechanism of Laa1 function in membrane traffic in budding yeast.

## Materials and Methods

The strains, plasmids, and oligonucleotides used for this study are listed in Tables 2-4 (James et al., 1996; Noguchi et al., 2008; Plemel et al., 2021). pFA6A-mcherry::His6Mx was generated by replacing GFP in pFA6a-GFP::His6Mx with mCherry from pRSETB-mcherry (Shaner et al., 2004). pMD397(pGAD-C1- Lft1) and pMD380 (pGEX4T1-Lft1) was generated by amplifying *LFT1* from genomic DNA using oligonucleotides FJ064 and FL065 and subcloning into pGADC1 between the BamHI and SalI sites. pMD394-396 were generated by amplifying from pMD380 with oligo pairs described in Table 4 and using 3-piece assembly reactions to generate the deletion alleles (Gibson et al., 2009). Tags and deletions were integrated into the genome using a one-step PCR-based method (Longtine et al., 1998; Noguchi et al., 2008). Mutations in *LFT1* were generated by a two-piece PCR-mediated gene replacement strategy, the *LFT1* wild-type or mutant open reading frames were amplified from pMD380 and pMD394-pMD396 using oligonucleotides MD1258 and MD1259 for wild-type and all mutations except the deletion of helix 1, which was amplified with MD1269b and MD1259. The open reading frame fragment was co-transformed with a fragment amplified from pFA6a-6His-TEV-Flag-His6Mx using MD902 and MD904 into cells carrying a deletion of *LFT1.* Combinations of tags, mutations, and deletions were performed using manual tetrad dissection. The functionality of Lft1-GFP was assessed by a sertraline sensitivity assay (Fig. S0).

Yeast cells were grown in yeast/peptone medium supplemented with 2% glucose and a mixture of adenine, uracil, and tryptophan (YPD+AUT), or yeast/peptone medium supplemented with 2% galactose and a mixture of adenine, uracil, and tryptophan (YPG+AUT) or synthetic medium SD supplemented with 2% glucose and an amino acid mix as previously described (Lang et al., 2014). Sertraline was from Fisher Scientific. Sertraline (10 mm) was prepared in DMSO. Antibodies against Myc (9E10, RRID:CVCL_L708) were from Biolegend, PGK1 (22C5D8,RRID:AB_2532235) was from Novex, HA (12CA5, RRID:AB_514505) and Flag (M2, RRID:AB_439685) were from Sigma. GST-antibody and Ent5 antibodies were described previously (Hung et al., 2012). Alexa Fluor 647-conjugated goat anti-mouse (RRID:AB_141698) and Alexa Fluor 647 goat anti-rabbit (RRID:AB_141663) antibody were from Life Technologies.

Graphs were generated and statistical analysis was performed with GraphPad Prism v7.00-9.00 (GraphPad Software, La Jolla, CA).

Imaging and analysis was performed as described previously (Hung and Duncan, 2016). Briefly, unless indicated otherwise cells were cultured to logarithmic phase in SD medium, centrifuged briefly to concentrate cells, and mounted on an uncoated coverslip in growth medium. Images were collected with a Nikon Ti-E inverted microscope with a 1.4 numerical aperture/100× oil immersion objective. The Lumencor LED light engine (472/20 nm for GFP and 575/20 nm for mCherry) was used for fluorophore excitation. Filters were ET/GFP-mCherry (59,002×), excitation dichroic (89019bs), and emission-side dichroic (T560lpxr), and for emission filters, ET525/50 m and ET595/50 m(Chroma).

The intensity of fluorescence puncta in cells was quantified as previously described (Martinez-Marquez and Duncan, 2018). Z-stack images were first compressed to a single maximum-intensity image using ImageJ. The resulting image was denoised using the “Denoise image” menu from the SpatTrackV2 software (Lund et al., 2014). Image analysis was performed in ImageJ. To define regions of GFP localization, the denoised image was thresholded using Otsu’s method with ImageJ with the stacked histogram option selected. GFP positive regions of interest were defined using the “Analyze Particles” menu from ImageJ on the binary thresholded image. The GFP-positive regions of interest were then used to measure fluorescence intensity from a background subtracted Z-stack compressed but non-denoised image. Fluorescence intensity was determined from at least eight fields of view per sample, each containing at least five cells. For plasma membrane to vacuole ratio analysis of Sna2^Y75A^, images were first background subtracted using a 50nm rolling ball in ImageJ, then linear regions of interest bisecting the plasma membrane or vacuolar membrane were defined. The ratio was calculated by dividing the maximum intensity value from the plasma membrane region of interest by the maximum intensity value from the vacuolar membrane region of interest. Experiments were repeated as indicated in the figures using at least two different strains for the same genotype.

For co-localization analysis, GFP and mCherry positive regions of interest were defined on denoised single plane images using SpatTrackV2 software (Lund et al., 2014). Image analysis was performed in ImageJ. To define regions of GFP or mCherry localization, the denoised image was thresholded using Otsu’s method with ImageJ with the stacked histogram option selected. Regions of interest were defined using the “Analyze Particles” menu from ImageJ on the binary thresholded image. The regions of interest were then used to overlap between GFP and mCherry signal. GFP structures that contain mCherry, were defined as structures for which at least 30% of the GFP positive area of the structure was also mCherry positive and vice versa for mCherry structures that contain GFP. At least eight fields of view were analyzed per sample, each containing at least five cells. Experiments were repeated as indicated in the figures using two different strains for each genotype.

For Drs2 localization, cells were transferred to a tube roller at 25C at least 3 hours before treating with 76 µg/ml Latrunculin or 2% DMSO. Cells were incubated for 30 min at 25C in microfuge tubes with punctured lids in a shaking heating block, before images were collected. Z-stack images were first compressed to a single maximum-intensity image using ImageJ. Files were copied and renamed to obscure genotype information and cells were manually scored as having punctate or cell surface localization by a single person who had established a scoring schema based on pilot data. At least five fields of view were analyzed per sample, each containing at least five cells.

For immunoblotting, after SDS-PAGE, samples were transferred to nitrocellulose, blocked with 5% milk in TBS-T (137 mm NaCl, 15.2 mm Tris-HCl, 4.54 mm Tris, 0.896 mm Tween 20), and then probed with primary and fluorescent secondary antibodies. Fluorescence signals were detected on an Azure 600 imaging system (Azure Biosciences). Image analysis was performed in ImageJ.

Whole-cell yeast extracts for protein abundance assessment were generated using glass bead lysis in SDS sample buffer as previously described (Aoh et al., 2011). Cell lysates for immunoprecipitation and GST- pulldowns were generated by glass-bead lysis in HEKG5 (20 mm Hepes, pH 7.5, 1 mm EDTA, 150 mm KCl, 5% glycerol) with protease inhibitor mixture without EDTA (Promega), followed by the addition of 0.3% CHAPS. The lysates were clarified by centrifugation at 13,000 rpm for 10 min at 4 °C. Protein concentrations were determined using Biorad Protein Assay (Biorad) and protein concentrations were adjusted to 2mg in 500 µl HEKG5. For Flag-IP, 6 µls of EZview™ Red ANTI-FLAG® M2 Affinity Gel (Sigma). For GST-pull downs, 20µls 50% Glutathione agarose (G-Biosciences) was added. For myc-IP, 20 µls 20% Protein A Sepharose (Cytvia) and 0.5 anti-myc antibodies were added. The lysates were incubated two hours at 4 °C. For immunoprecipitations, the beads were washed three times with ice-cold HEKG5 with 0.3% CHAPS, and the bound proteins were eluted with SDS sample buffer. For GST- pulldowns, the beads were wash twice with HEKG5 with 0.3% CHAPS and once with HEKG5 without CHAPS. The beads were resuspended in 50 µl 100 mm Tris, pH 9, 200 mm NaCl, 5 mm DTT, 20 mm reduced GSH, and incubated for 10 min at room temperature. The supernatants were reserved. A second elution was then performed the two supernatants were combined. For all samples, cells were cultured to mid-log phase in YPD+AUT except for samples described in Figure 6 A and Figure 7 A which were grown in YPG+AUT.

For mass spectrometry analysis, yeast lysates were prepared by liquid nitrogen lysis (Goode, 2002). Frozen powder was thawed into room temperature HEKG5 with 0.3% CHAPS and protease inhibitor mixture without EDTA. The lysates were clarified by centrifugation at 13,000 rpm for 10 min at 4 °C. Protein concentrations were determined using Biorad Protein Assay and protein concentrations were adjusted to 3.5mg in 1ml HEKG5. 40 µl of GFP-TRAP beads (Chromotek) beads were added and samples were incubated for 30 minutes at 4°C. Beads were washed 3 times with HEKG5 with 0.3% CHAPS and protease inhibitor mixture without EDTA and washed 3 times with 20 mm HEPES, pH 7.5, 1 mm EDTA, 150 mm KCl. The beads were aspirated dry, snap frozen, and stored at -80°C.

Approximately the sample was submitted to Proteomics Resource Facility (PRF) for high-resolution LC- MS/MS analysis. Analysis was performed per protocol optimized at the PRF. Briefly, cysteines were reduced with 10 mM DTT (45° C for 30 min), and alkylation of cysteines was achieved by incubating with 50 mM 2-Chloroacetamide, under darkness, for 30 min at room temperature. An overnight digestion with 0.5 ug sequencing grade, modified trypsin (Promega) was carried out at 37°C with constant shaking in a Thermomixer. Digestion was stopped by acidification and peptides were desalted using SepPak C18 cartridges using manufacturer’s protocol (Waters). Samples were completely dried using vacufuge. Resulting peptides were dissolved in 20 µl of 0.1% formic acid/2% acetonitrile solution and 2 µls of the peptide solution were resolved on a nano-capillary reverse phase column (Acclaim PepMap C18, 2 micron, 50 cm, ThermoScientific) using a 0.1% formic acid/2% acetonitrile (Buffer A) and 0.1% formic acid/95% acetonitrile (Buffer B) gradient at 300 nl/min over a period of 90 min (2-25% buffer B in 45 min, 25-40% in 5 min, 40-90% in 5 min followed by holding at 90% buffer B for 5 min and requilibration with Buffer A for 30 min). Eluent was directly introduced into Q exactive HF mass spectrometer (Thermo Scientific, San Jose CA) using an EasySpray source. MS1 scans were acquired at 60K resolution (AGC target=3x106; max IT=50 ms). Data-dependent collision induced dissociation MS/MS spectra were acquired using Top speed method (3 seconds) following each MS1 scan (NCE ∼28%; 15K resolution; AGC target 1x105; max IT 45 ms).

Proteins were identified by searching the MS/MS data against S cerevisiae protein database (UniProt; 5953 entries; downloaded on 01/22/2020) using Proteome Discoverer (v2.4, Thermo Scientific). Search parameters included MS1 mass tolerance of 10 ppm and fragment tolerance of 0.1 Da; two missed cleavages were allowed; carbamidimethylation of cysteine was considered fixed modification and oxidation of methionine, deamidation of asparagine and glutamine were considered as potential modifications. False discovery rate (FDR) was determined using Percolator and proteins/peptides with a FDR of ≤1% were retained for further analysis.

Two-hybrid interactions were monitored as previously described (Duncan et al., 2003). IC50 values were determined as previously described (Hung et al., 2018).

Vacuole labeling with FM4-64 (Synaptored^TM^ C2 from Biotium) was performed as previously described (Buelto et al., 2020).

Structure predictions were performed using alphafold2_multimer_v3 version 1.5.2 with the following settings: mas_mode mmseqs2_uniref_env, 5 models, 6 recycles, 1 ensemble (Mirdita et al., 2022)

Structure comparisons and alignments were performed in UCSF ChimeraX version 1.4 with the following settings: Sequence alignment algorithm Needleman-Wunsch, Matrix Blosum-62, Gap extension penalty 1, iterate by pruning long atom pairs was selected and cutoff distance was set at 2 (Goddard et al., 2018; Pettersen et al., 2021).

## Online Supplemental material

Table S1 contains mass-spectrometry results. Table S2 contains a list of used. Table S3 contains a list of plasmids used. Table S3 contains a list of oligonucleotides used. Figures S1 and S2 show full structure predictions, and PAE plots for Laa1-Lft1 and Laa1-Laa2 complexes. Figure S3 shows GFP-NSI assay data showing no effect of loss of Lft1, Laa2, or Laa1 on AP3 mediated traffic. Figure S4 shows immunoprecipitation data showing no detectable interaction between Ent5 and Lft1, Laa2, or Laa1. Figure S5-8 shows full structure predictions, and PAE plots for Heatr5A-Aftiphilin and HeatrA-Clba1complexes. Figure S9 shows sertraline sensitivity data indicating functionality of Lft1-GFP when expressed at the endogenous locus.

## Acknowledgments

UCSF ChimeraX was developed by the Resource for Biocomputing, Visualization, and Informatics at the University of California, San Francisco, with support from the National Institutes of Health R01- GM129325 and the Office of Cyber Infrastructure and Computational Biology, National Institute of Allergy and Infectious Diseases. We acknowledge support from the University of Michigan Biomedical Research Core Facilities (Proteomics Resource Facility).

This work was supported by the National Institutes of Health (R01-GM129255, UL1-TR002240). Microscopy was performed on a system generously provided by Ajit P. Joglekar. We thank Ming Li, Yanzhuang Wang and Ajit Joglekar for helpful comments on this manuscript. We thank the anonymous reviewers for helpful comments, and for highlighting the importance of the Whitfield dissertation to this study.

## Author contributions

L.J. Marmorale, H. Jin, T. Reidy, and M.C. Duncan conceived the experiments. L.J. Marmorale, H. Jin, T. Reidy, C. Zysnarski, F. Jordan-Javed, and S. Lahiri generated the reagents. L.J. Marmorale, H. Jin, T. Reidy, C. Zysnarski, B. Palomino-Alonso, and M.C. Duncan performed the experiments and analyzed the data. M.C. Duncan wrote the manuscript and secured funding.

---

**Figure S1.**
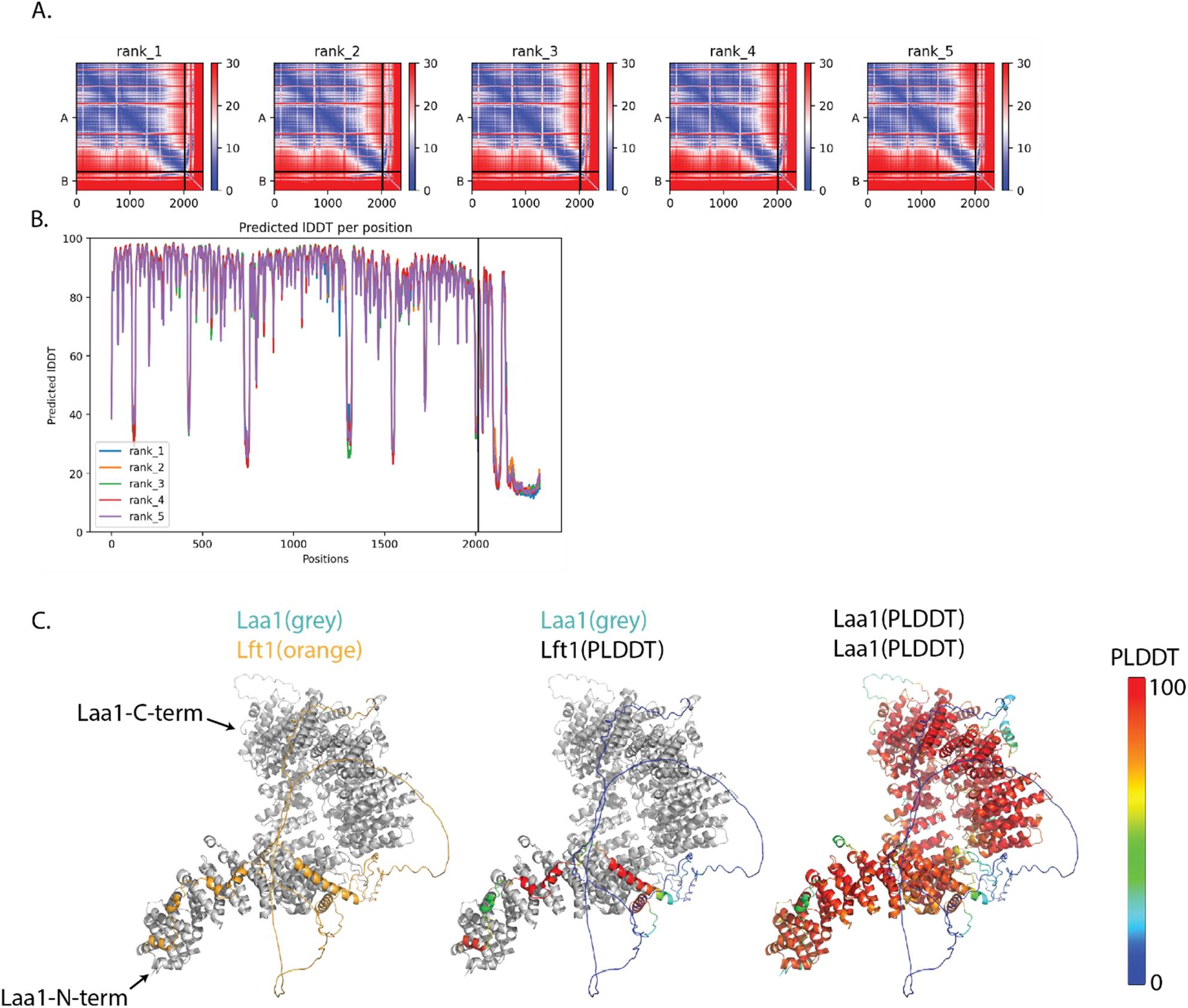
Structure prediction of Laa1 interaction with Lft1. A. Predicted Alignment Error plots for top 5 ranked models with Lft1 as chain B. B. Predicted local distance difference test for top 5 ranked models. C. Rank 1 model. Left panel shows Lft1 in orange and Laa1 in grey. Center panel shows Lflt1 colored by PLDDT values and Laa1 in grey. Right show both proteins colored by PLDDT.

**Figure S2.**
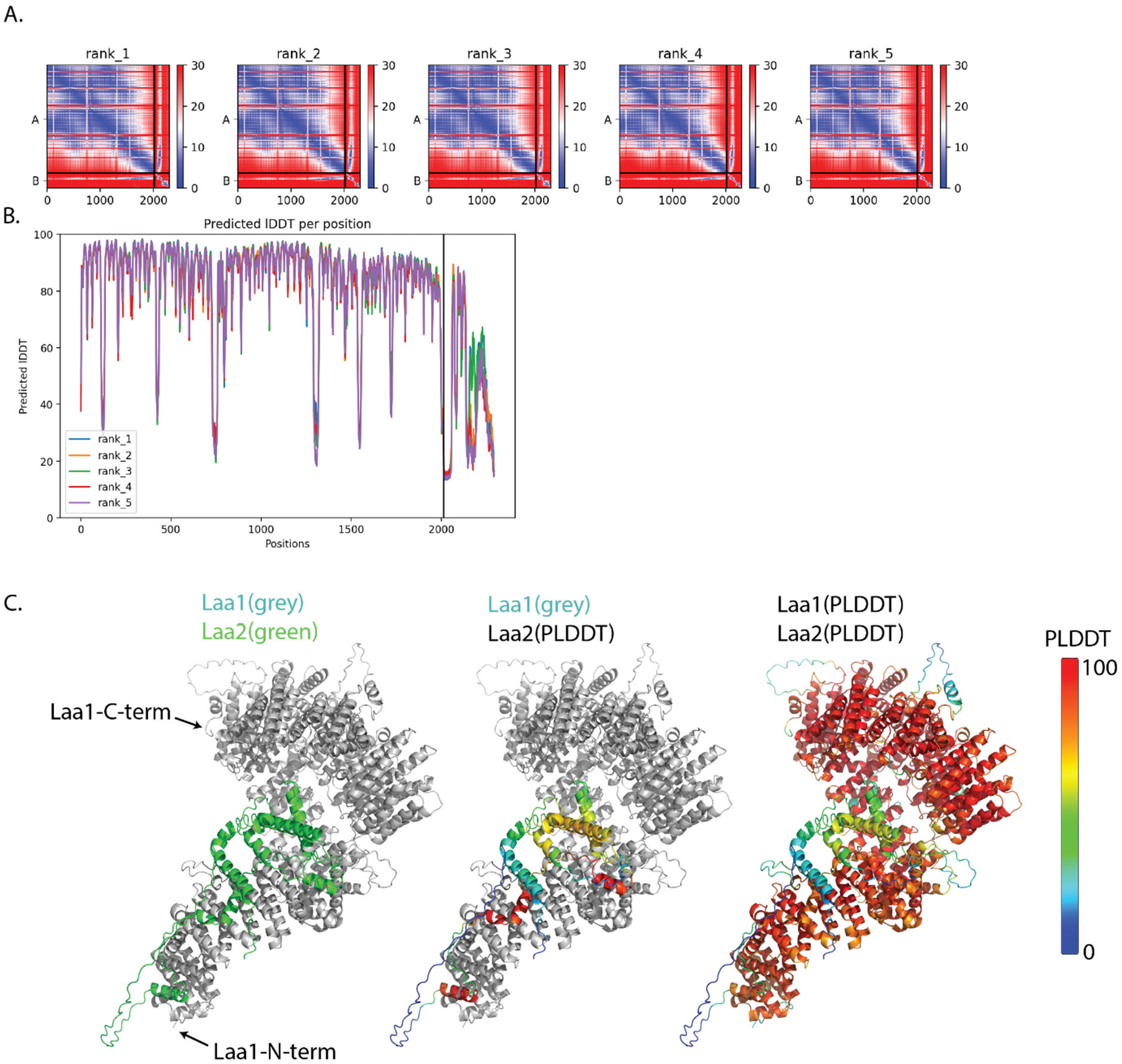
Structure prediction of Laa1 interaction with Laa2. A. Predicted Alignment Error plots for top 5 ranked models with Laa2 as chain B. B. Predicted local distance difference test for top 5 ranked models. C. Rank 1 model. Left panel shows Laa2 in green and Laa1 in grey. Center panel shows Laa2 colored by PLDDT values and Laa1 in grey. Right show both proteins colored by PLDDT.

**Figure S3.**
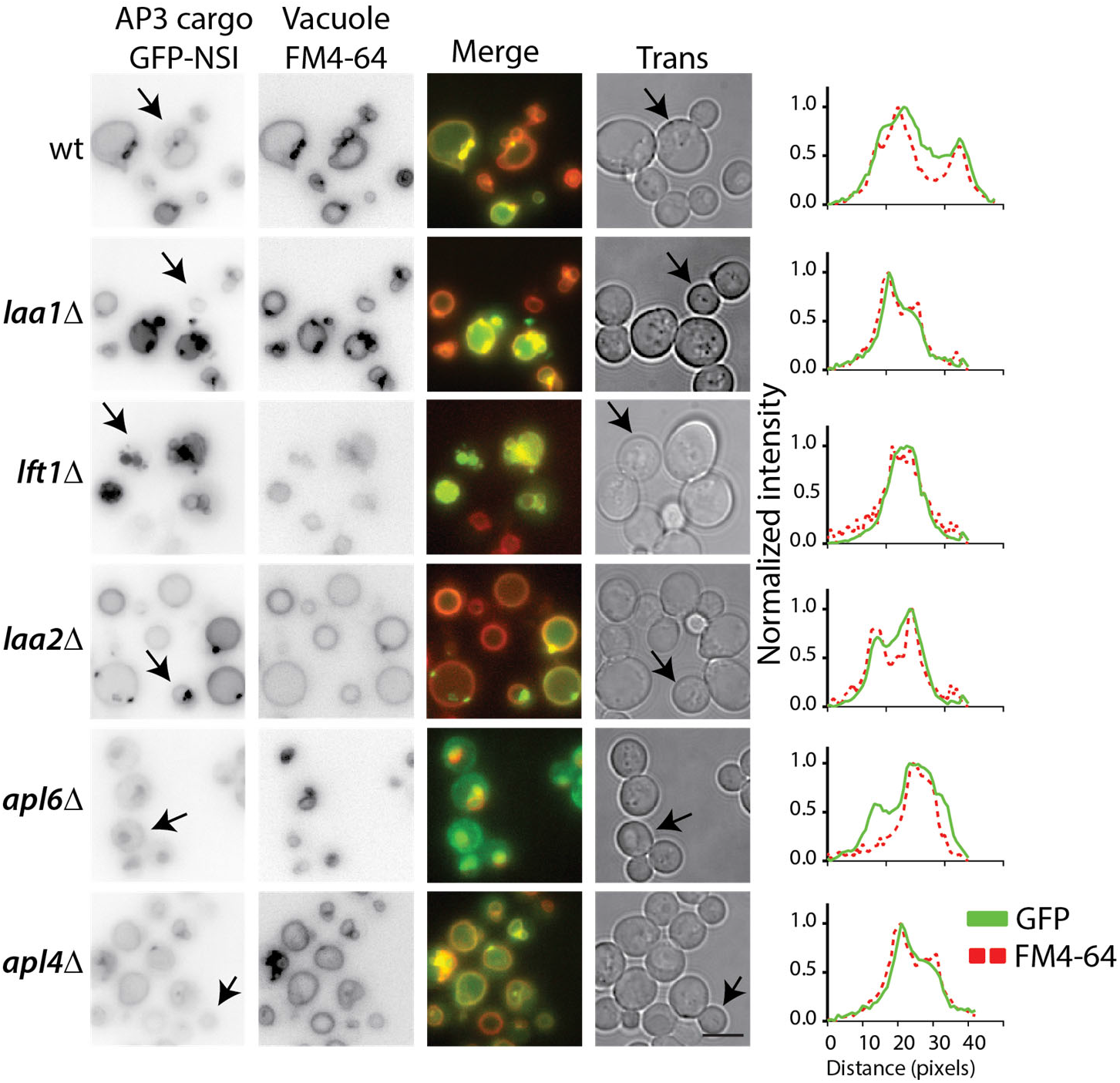
The AP3 pathway is not disrupted in cells lacking Laa1, Lft1, Laa2, or Apl4. Micrographs of indicated genotypes expressing GFP-NSI, an AP3 cargo. Vacuoles were labeled by incubation with FM4-64 for 20 mins, excess dye was washed, and FM4-64 was chased into the vacuole for 30 mins before imaging. In wild-type cells and most mutants tested, GFP is found exclusively in the vacuole. When AP3 (apl6D) is disrupted the GFP signal is observed at the plasma membrane and vacuole. Arrows show cell periphery identified from transmitted light image overlaid on GFP image. Charts show line scans of the GFP and FM4-64 signal corresponding to the crosssection indicated by the arrow, data are normalized to min and max values of each line scan. Bar is 5um.

**Figure S4.**
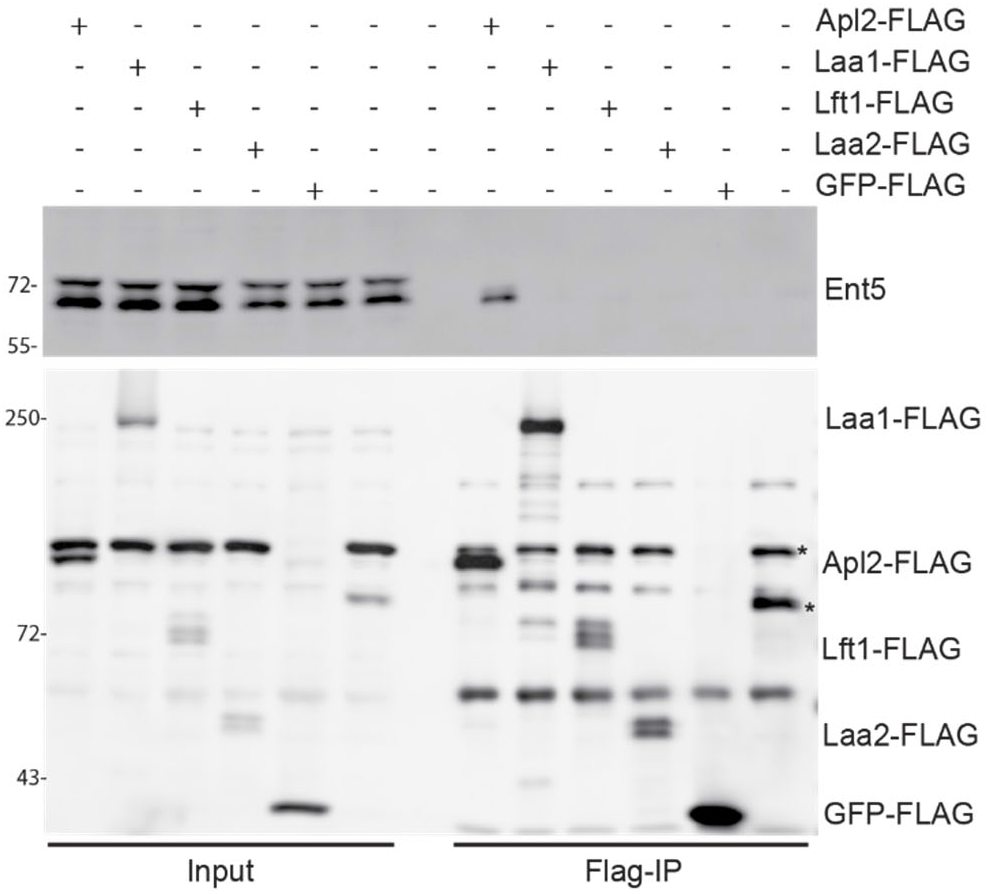
Lft1 does not detectably bind to Ent5. Ent5 was not detected in coimmunoprecipitation samples with Laa1, Lft1, or Laa2 under conditions where Ent5 binds AP1. Flag-proteins were immunoprecipitated from cells expressing indicated gene fusions from their endogenous loci and probed for Ent5 and Flag. * indicates flag crossreacting band found in the BY background. The GFP-Flag strain comes from a cross to FY5, the parent of the BY background, which lacks this band.

**Figure S5.**
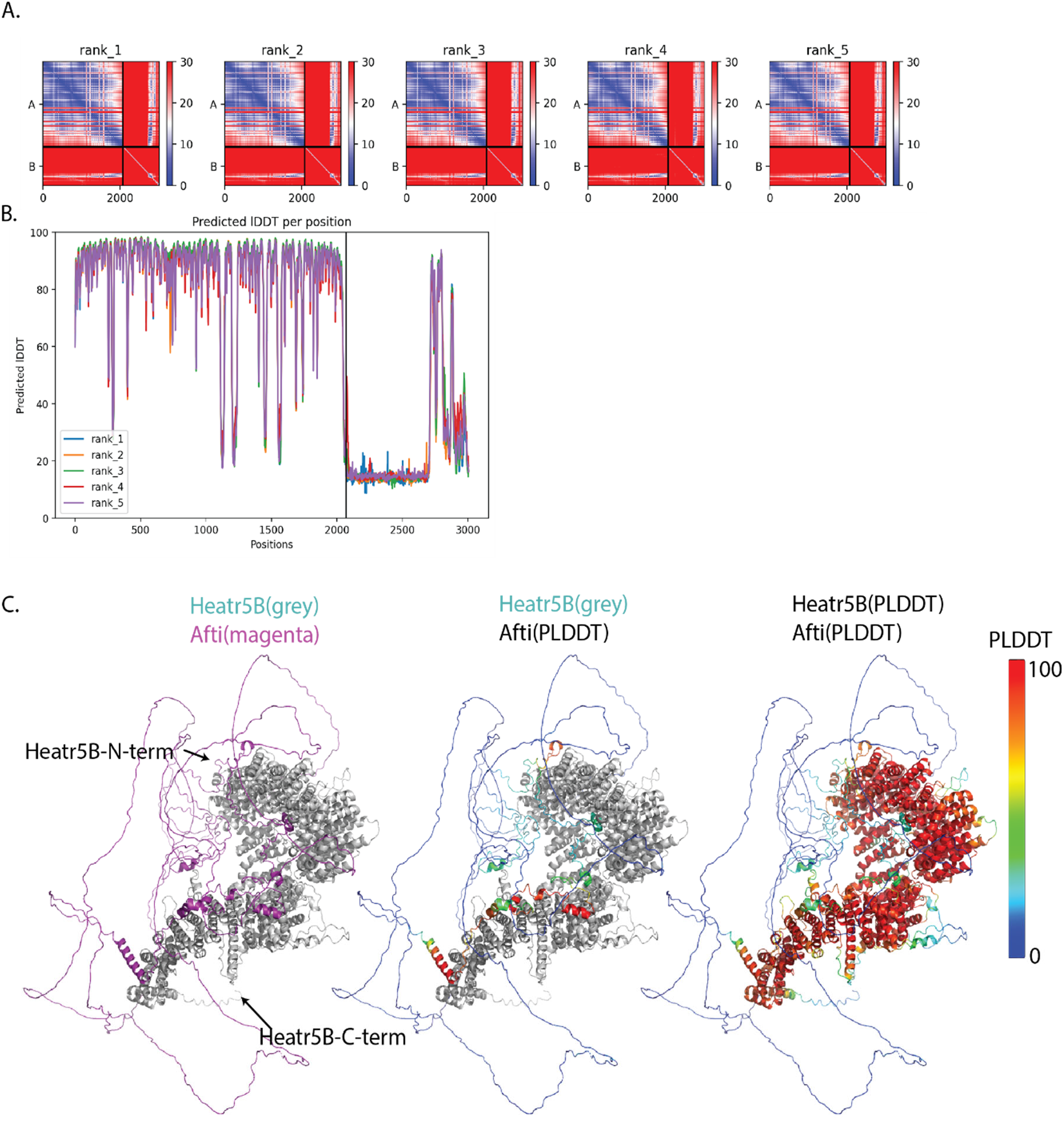
Structure prediction of Heatr5B interaction with Aftiphilin. A. Predicted Alignment Error plots for top 5 ranked models with Aftiphilin as chain B. B. Predicted local distance difference test for top 5 ranked models. C. Rank 1 model. Left panel shows Aftiphilin in magenta and Heatr5b in grey. Center panel shows aftiphilin colored by PLDDT values and Heatr5b in grey. Right show both proteins colored by PLDDT.

**Figure S6.**
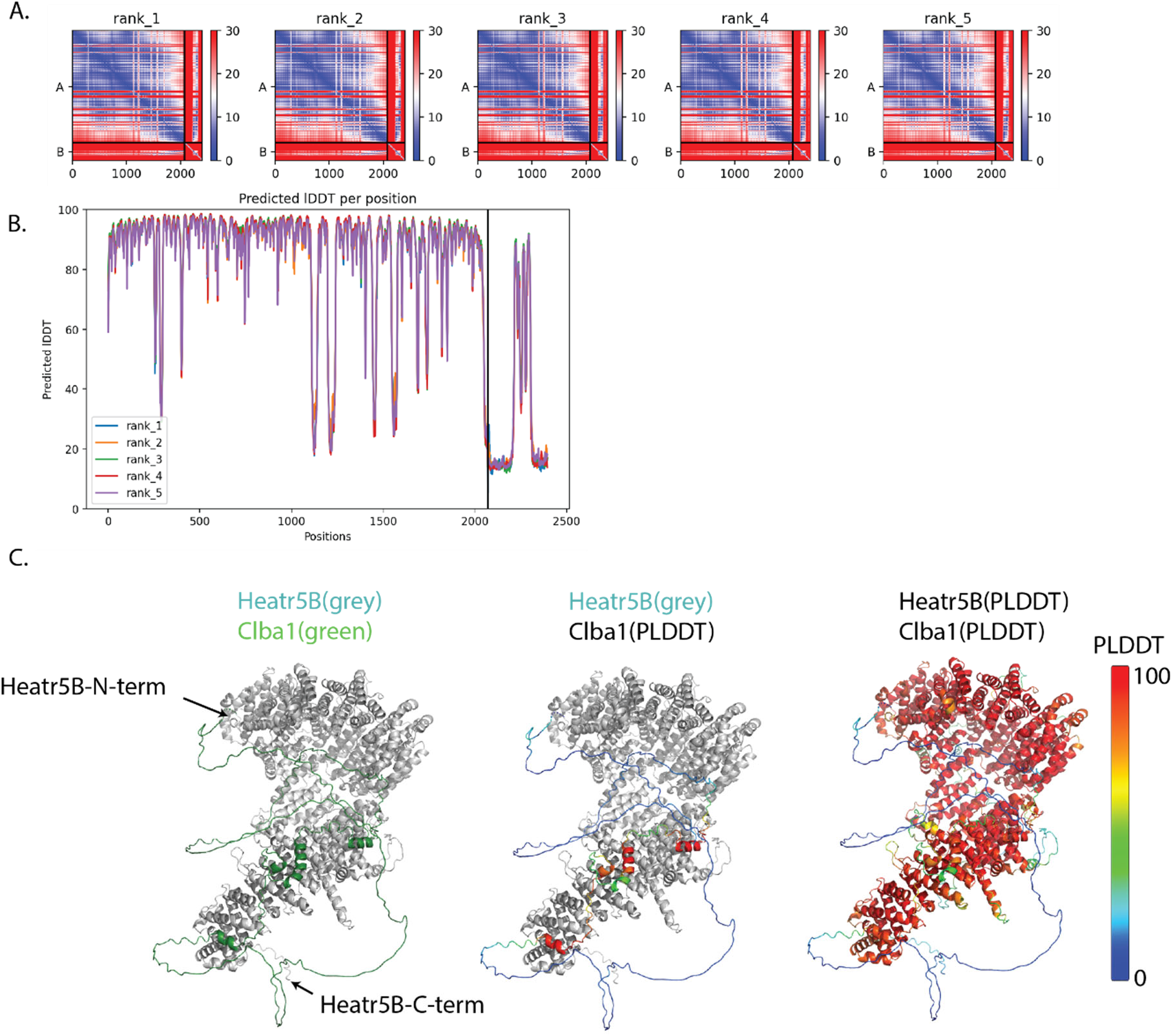
Structure prediction of Heatr5B interaction with Clba1. A. Predicted Alignment Error plots for top 5 ranked models with Clba1 as chain B. B. Predicted local distance difference test for top 5 ranked models. C. Rank 1 model. Left panel shows Clba1 in green and Heatr5b in grey. Center panel shows Clba1 colored by PLDDT values and Heatr5b in grey. Right show both proteins colored by PLDDT.

**Figure S7.**
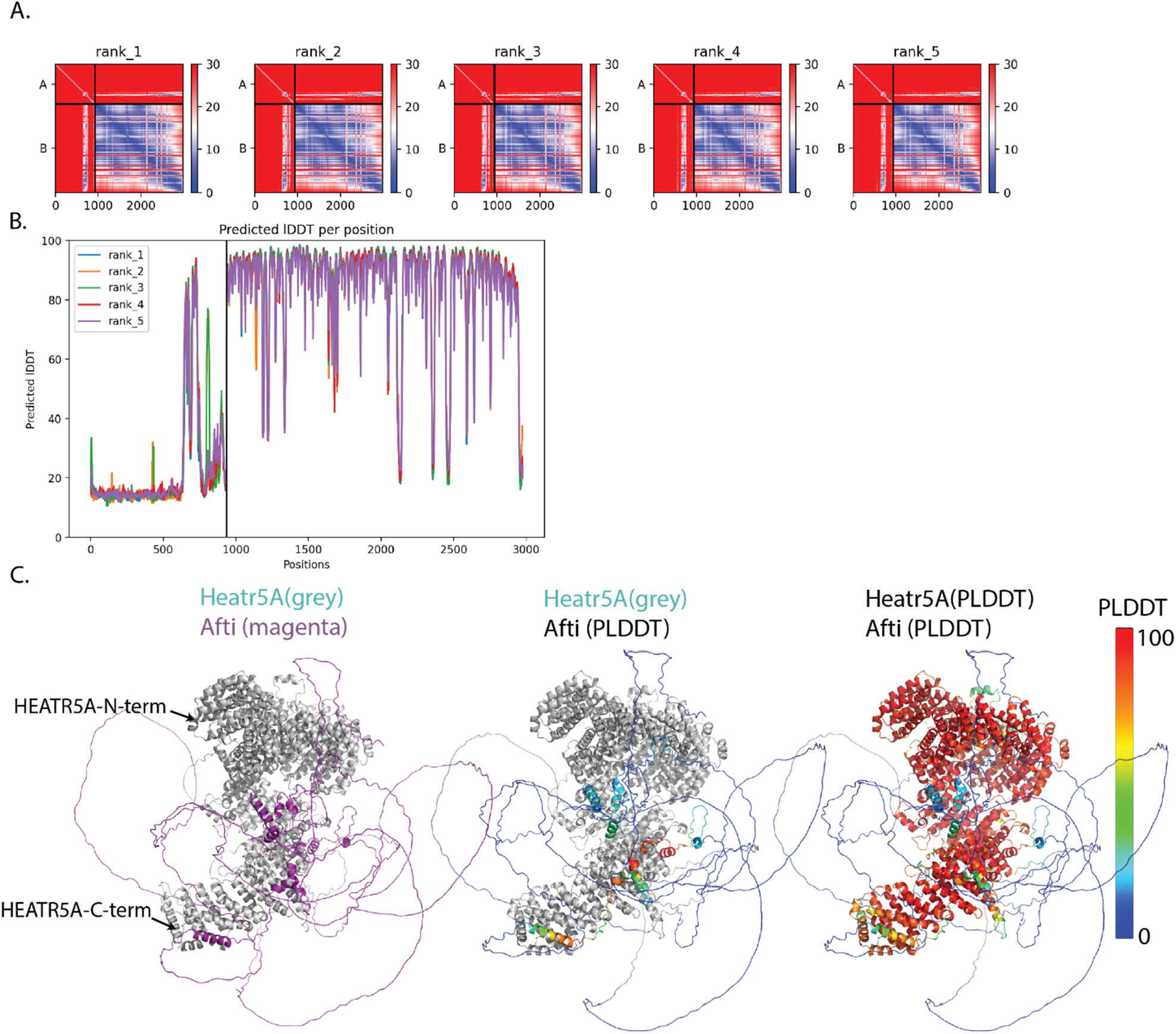
Structure prediction of Heatr5a interaction with Aftiphilin. A. Predicted Alignment Error plots for top 5 ranked models with Aftiphilin as chain A. B. Predicted local distance difference test for top 5 ranked models. C. Rank 1 model. Left panel shows Afti in magenta and Heatr5a in grey. Center panel shows Aftiphilin colored by PLDDT values and Heatr5a in grey. Right show both proteins colored by PLDDT.

**Figure S8.**
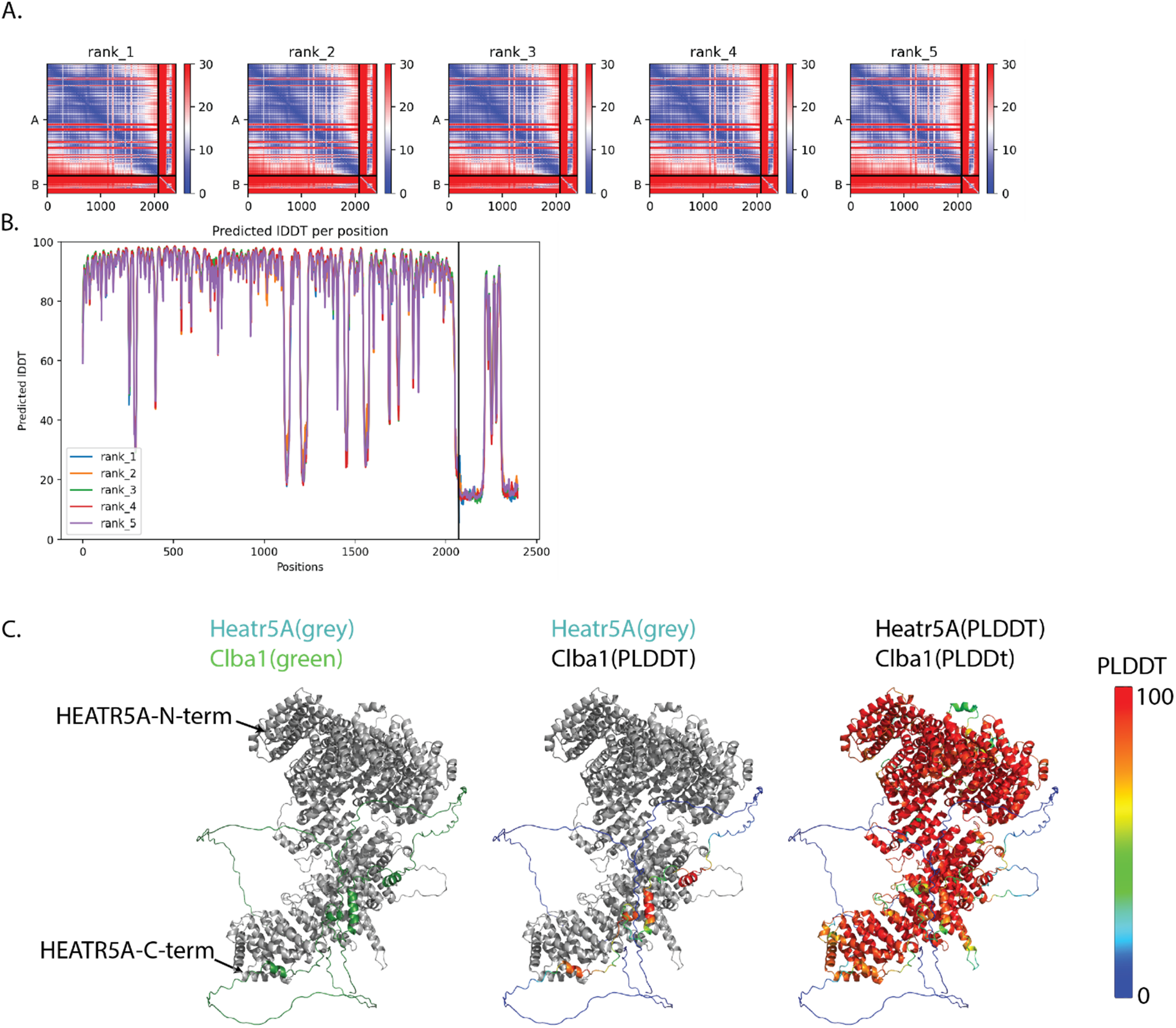
Structure prediction of Heatr5a interaction with Clba1. A. Predicted Alignment Error plots for top 5 ranked models with Clba1 as chain B. Predicted local distance difference test for top 5 ranked models. C. Rank 1 model. Left panel shows Clba1 in green and Heatr5a in grey. Center panel shows Clba1 colored by PLDDT values and Heatr5a in grey. Right show both proteins colored by PLDDT.

**Figure S9.**
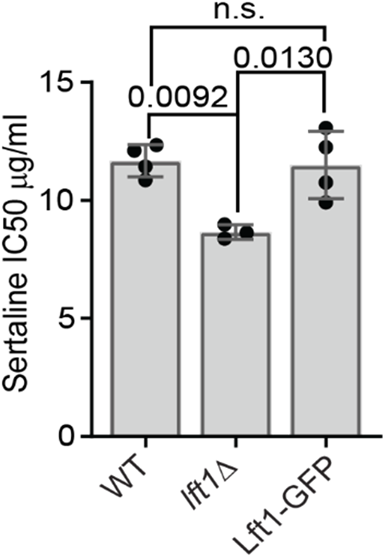
Lft1-GFP does not increase sensitivity to sertraline. To test the functionality of Lft1-GFP, the sensitivity of cells expressing Lft1-GFP to sertraline was determined. Chart shows IC50s from replicate experiments, mean and standard deviation. P-values were calculated with Tukey’s multiple comparisons test.

